# LeafGo: Leaf to Genome, a quick workflow to produce high-quality *De novo* genomes with Third Generation Sequencing technology

**DOI:** 10.1101/2021.01.25.428044

**Authors:** Patrick Driguez, Salim Bougouffa, Karen Carty, Alexander Putra, Kamel Jabbari, Muppala Reddy, Richard Soppe, Nicole Cheung, Yoshinori Fukasawa, Luca Ermini

## Abstract

Recent years have witnessed a rapid development of sequencing technologies. Fundamental differences and limitations among various platforms impact the time, the cost and the accuracy for sequencing whole genomes. Here we designed a complete *de novo* plant genome generation workflow that starts from plant tissue samples and produces high-quality draft genomes with relatively modest laboratory and bioinformatic resources within seven days. To optimize our workflow we selected different species of plants which were used to extract high molecular weight DNA, to make PacBio and ONT libraries for sequencing with the Sequel I, Sequel II and GridION platforms. We assembled high-quality draft genomes of two different *Eucalyptus* species *E. rudis*, and *E. camaldulensis* to chromosome level without using additional scaffolding technologies. For the rapid production of *de novo* genome assembly of plant species we showed that our DNA extraction protocol followed by PacBio high fidelity sequencing, and assembly with new generation assemblers such as hifiasm produce excellent results. Our findings will be a valuable benchmark for groups planning wet- and dry-lab plant genomics research and for high throughput plant genomics initiatives.

## Introduction

Plants represent the dominant kingdom of life in terms of Earth biomass [1], and through colonization of terrestrial and aquatic habitats, are responsible for maintaining ecological and atmospheric balance. Despite being globally distributed, climate change and anthropogenic activities are massively impacting current plant diversity with repercussions for ecophysiology, distribution and interactions with other organisms [2,3]. Sequencing present-day plant genomes to better understand genomic diversity is an important requirement for gauging plants’ susceptibility to climate change.

In the last decade, rapid advances in short read sequencing technology has resulted in the availability of over 300 plant species genomes, of differing quality [4]. Recently, long-read sequencing methods (Pacific Biosciences, PacBio, and Oxford Nanopore Technology, ONT) are becoming more accessible while technological advances have led to increases in the base accuracy and the sequencing length, as well as a significant reduction in cost per base of sequence [5]. The main benefit of long-read sequencing technologies for genomics, compared to the more dominant short-read/Illumina sequencing, is the ability to assemble genomes relatively easily by linking reads that span across repetitive genomic regions. This property when combined with ultra-long reads, highly accurate sequencing and complementary scaffolding technologies has thereby enabled highly accurate telomere to telomere assemblies [6–8]. The many benefits of long-read sequencing of large DNA fragments has boosted demand for high quality high molecular weight (HMW) DNA.

In the case of plants, genomes can be large and highly repetitive making long-read sequencing ideal for genome assembly [9–11]. However, it is often difficult to extract HMW DNA suitable for long-read sequencing from plants [12–14]. Plants have tough cell walls and contain high levels of metabolic contaminants, such as polyphenols and polysaccharides [15–17] which are difficult to eliminate and impact sequencing quality. Furthermore, a plethora of new initiatives have emerged to sequence millions of species, including those from the plant kingdom [18,19]. To address the need to produce high quality genomes for thousands of plant species we developed a workflow (Fig. 1) that efficiently produces HMW DNA suitable for long-read sequencing and results in data that can be rapidly assembled into a genome with relatively modest resources within seven days. To test our workflow for HMW DNA extraction we selected seven species of plant (*Salicornia bigelovii*, *Distichlis palmeri*, *Eucalyptus rudis* subsp. *rudis*, *E. camaldulensis* subsp. *obtusa*, *Salvadora persica*, *Zea mays*, and *Pennisetum glaucum*) from diverse taxonomic groups. Maize (*Z. mays*) and pearl millet (*P. glaucum*) are well-studied, globally-important crops responsible for feeding millions of people with large, highly repetitive genomes [10,20–24]. Nipa grass (*D. palmeri*), dwarf saltwort (*S. bigelovii*) and toothbrush tree (*S. persica*) are lesser-researched plants without published genomes that are, or could be developed into, agriculturally/pharmacologically important crops [25–28]. Finally, the flooded gum (*E. rudis*) and river red gum (*E. camaldulensis*) trees were selected. Eucalypts are the most commonly planted hardwood trees in the world due to their fast growth, environmental adaptability and many commercial uses [29]. Good quality HMW DNA is relatively difficult to extract from *Eucalyptus* species due to their high phenolic and polysaccharide content [12,17,30]. HMW DNA was processed into long read sequencing libraries for the PacBio and ONT GridION platforms, and the data quality assessed for downstream analysis. In particular, PacBio Continuous Long Read (CLR) and High Fidelity (HiFi, also known as circular consensus sequencing, CCS) libraries were sequenced on the latest PacBio platforms, Sequel I and Sequel II. Finally, the two *Eucalyptus* species were assembled into novel high-quality draft genomes using the latest tools.

**Figure 1:**
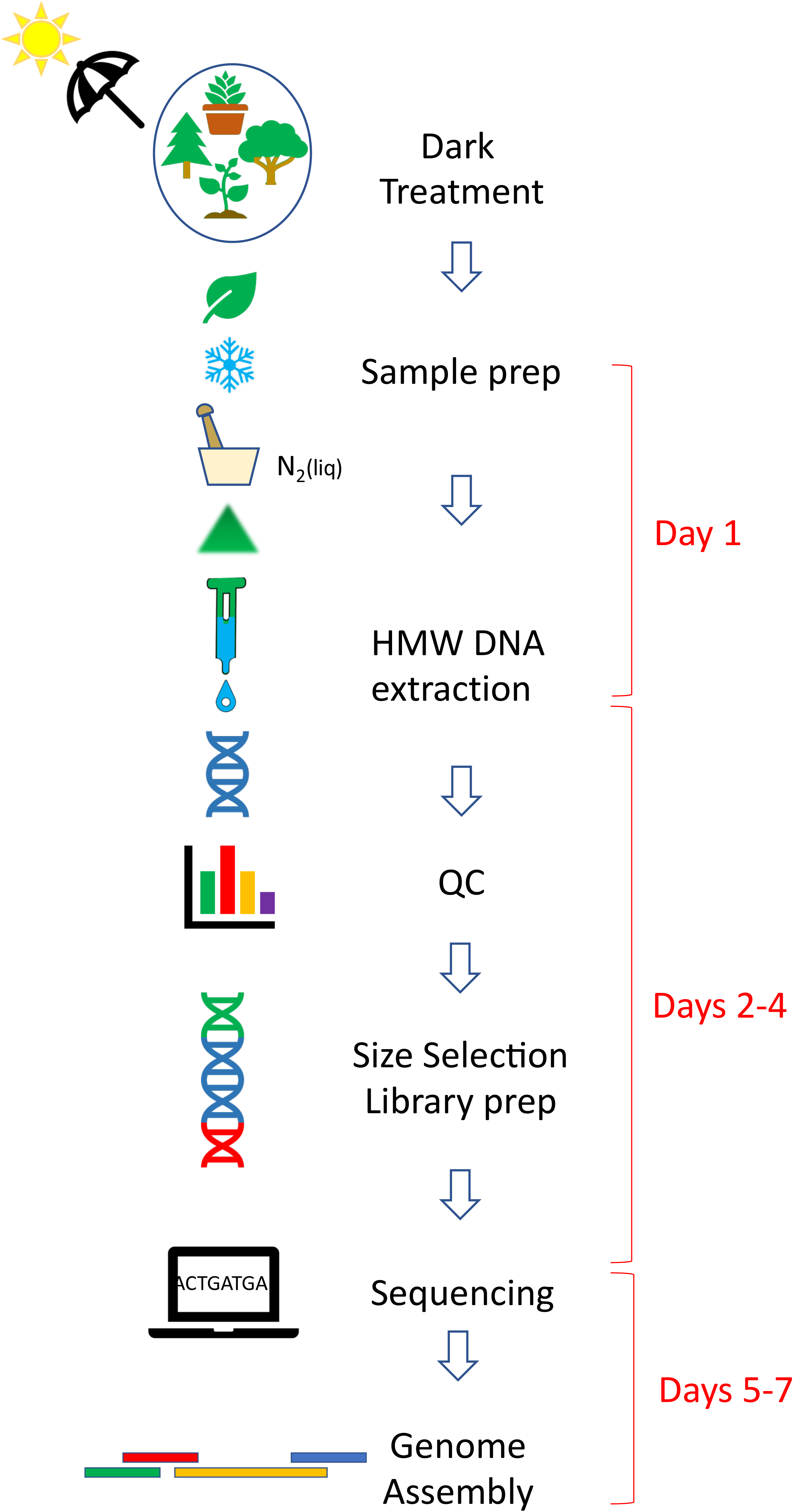
Plant Long-Read Sequencing (LeafGo) Workflow

## Results

### DNA extraction, quality controls, library and long read sequencing

#### DNA extraction

Our extraction protocol generated high pure and copious amounts of HMW DNA within a day and using minimal resources and effort (Supp. Fig. 1, 2 and Supp. Table 1). The protocol reproducibly yielded high quality HMW DNA in 20 separate extractions from seven different plant species over different days by different technicians. The yield (per 1 g wet weight of leaf) ranged from 15 to 278 μg with high variability in the *Eucalyptus* species, compared to the other species (Supp. Table 1). All the extracted HMW DNA had high purity and integrity, despite the different composition, including potential contaminants, between species [12,25,28,30,31]. Notably, the *Eucalyptus* samples showed signs of oxidation during lysis, such as dark colouration of the solution; however, the quality of DNA was not compromised, without the need for added antioxidants [12,30]. The absorbance ratios (Supp. Table 1) indicated low levels of contaminants, such as protein, carbohydrates and phenolics, and high purity (A_260/280_ ~ 1.8; A_260/230_ > 2.1) [9,30–32]. The integrity of the extracted genomic DNA (gDNA) was assessed by performing pulsed-field gel electrophoresis and capillary electrophoresis. The extracted samples typically had a DNA smear ranging from approximately >15 kb to <150 kb and a modeof over 80 Kb for all samples (Supp. Fig. 1 and 2).

Our results are comparable to, or improve on, similar studies in terms of DNA yield, fragment length and purity but with a simpler and less toxic extraction protocol [12,17,30] that produces HMW gDNA suitable for long-read sequencing in one day.

#### PacBio CLR and HiFi sequencing

The extracted HMW gDNA from the seven species, was processed into 26 CLR and 5 HiFi libraries prior to sequencing with the PacBio Sequel I (22: 1M SMRT cells) and Sequel II (9: 8M SMRT cells). The CLR libraries (Supp. Fig. 3) showed a mode > 30 kb with few shorter fragments. The sequencing results demonstrate that the extracted gDNA was of good purity and of high molecular weight. The sequencing metrics were above PacBio recommended specifications [33] and the internal controls yielded optimal results indicating that no inhibition of the sequencing reaction was observed as shown by SMRT Link statistics (Supp. Table 2). The average CLR throughput per Sequel I SMRT cell was 11.5 gigabases (Gb) (min: 3.3 Gb; max: 18.2 Gb) while the Sequel II yielded 167.6 Gb/SMRT cell (140.8 - 195.4 Gb). The average N50 subread length was 36.6 kilobases (Kb, min: 21.5 Kb, max: 44.7 Kb) for the 22 Sequel I CLR libraries, and 36.4 Kb (32.4 - 39.8 Kb) for the four Sequel II CLR libraries (Supp. Table 2, Supp. Fig. 4). Sheared DNA was tested for CLR library preparation; however, there was no obvious improvement in yield or N50, as previously shown with ONT libraries [30], and unsheared DNA was used for all other libraries. To test whether library loading affected the subread N50 length some SMRT cells were intentionally underloaded. There was a borderline significant correlation between ZMW (zero mode waveguide) occupancy or library underloading (high P0%) and subread N50 (Spearman’s rho = 0.42, p = 0.05; Supp. Table 3 and Supp. Fig. 5) suggesting a small benefit in subread N50 at the expense of throughput yield; unsurprisingly, library loading (determined from P0% and P1%) was strongly correlated with throughput yield (Spearman’s rho = −0.84 & 0.86, p < 0.01). Although not presented here, initial testing with higher size selected CLR libraries (35 Kb and 40 Kb) yielded diminished sequencing results with inconsistent throughput and lower N50 sizes. For this reason, we exclusively used 30 Kb size-selected CLR libraries. The reason for the suboptimal results with higher size selected libraries is not clear and might require follow up studies.

The HiFi libraries showed a mode of approximately 20 Kb (Supp. Fig. 6) and, sequenced with Sequel II, yielded an average total throughput of 344.0 Gb/SMRT cell (113.9 - 477.3 Gb), with the Q20 yield of 22.2 Gb (7.6 - 29.4 Gb) and subread N50 of 19.8 Kb (17.6 - 22.2 Kb). The total throughput and Q20 yield was high for all the samples except for one *D. palmeri* HiFi library that was underloaded (P1% = 13%); subsequent resequencing of the library with optimal loading yielded more typical results (356.3 Gb total throughput; 23 Gb Q20 yield). Results are shown in Supp. Table 2.

#### Oxford Nanopore Technology Sequencing

In order to evaluate the suitability of our workflow with another long read technology, we sequenced the two *Eucalyptus* species from our test plant species set with ONT. This technology, unlike PacBio sequencing, has theoretically no upper limit on the maximum read length and thus the N50 parameter takes on particular importance. Previous plant studies showed the impact of size selection on the N50 [30,34], and thus we tested two different size-selection methods: a chemical precipitation approach (Short Read Eliminator, SRE-XL) and a gel-based method (BluePippin with 30 Kb cut) using HMW DNA extracted from the two *Eucalyptus* species. Results showed both methods equally able to reduce the amount of smaller DNA fragments and their comparability (Supp. Fig. 7). However, despite the precipitation-based approach requiring less lab time, the fragment size distribution is more spread therefore retaining more shorter fragments.

The sequencing N50 for all libraries was very similar (average 41.4 Kb, range: 37.2 Kb - 43.8 Kb) with no apparent difference between species or size-selection method (Supp. Table 4).

### *De novo* genome assembly of *Eucalyptus* species

To assess the impact of the library type on the genome assembly, we used the data of two *Eucalyptus* species in our test plant species set. Prior to assembly the raw sequencing data was quality checked and the genomic data was characterized. Finally, we assembled both PacBio CLR and HiFi sequencing data and determined the contiguity of the assemblies.

#### LongQC quality control of raw data

The basic sequencing parameters of each sequencing run were examined using SMRT Link for PacBio. A more comprehensive quality control step was performed on the raw sequencing data using LongQC (Fukasawa et al. 2020), a specialized QC package for long read data. HiFi data for both *Eucalyptus* species were good quality as indicated by high scores for per read base calling accuracy (Supp. Fig. 8A and 8F) and by a normal distribution of per read coverage, except for slight bimodality in *E. rudis* (Supp. Fig. 8B). Both *Eucalyptus* species showed a similar GC content and sequence complexity (Supp. Fig. 8C, 8H and 8D, 8I). The GC content for all the reads shows a sharp unimodal distribution around a mean of 0.39 (± standard deviation ± 0.03), but with an upper sub-mode outlier near 0.55. A closer inspection of this higher GC content peak revealed the presence of telomere repeats. No artificial sequence adapters are present in the flanking region for either datasets (Supp. Fig. 8E and 8L).

In contrast, CLR data does not achieve the same level of quality shown by HiFi data. The Phred scores are not provided by the Sequel platforms and thus read base calling accuracy cannot be directly assessed. The sequence complexity of CLR is lower (low complexity fraction: 0-0.4 and 0-0.2 in CLR and HiFi, respectively), and the flanking regions seem to extend over a hundred bases pointing to the possibility of artificial sequences (Supp. Fig. 9C, 9G and 9D, 9H). Both *Eucalyptus* species’ CLR sequences have a similar GC content to that for HiFi data. The GC content for all the reads has a sharp normal distribution around a mean of 0.4. (± 0.04), but the upper sub-mode peak, indicative of telomere repeats and present in HiFi data, is not found within the CLR distribution. Surprisingly, CLR sequencing may be less sensitive at detecting telomere repeats than HiFi.

#### Genome profiling

We used the software package GenomeScope 2.0 [36] to infer the size and the heterozygosity level of the two *Eucalyptus* genomes from unassembled HiFi sequenced data. The rationale is that if the genome is heterozygous, then the *k*-mer profile will exhibit a characteristic bimodal distribution, as shown in HiFi genome sequencing data (Supp. Fig. 10). We observed two peaks that were centered at coverage values of 20x and 40x in the case of *E. rudis* and 25x and 50x in the case of *E. camaldulensis*. These values represent both heterozygous *k*-mers that have been sequenced at half the coverage (20x and 25x coverage for *E. rudis* and *E. camaldulensis,* respectively) and homozygous *k-*mers sequenced equally from both alleles (40x and 50x coverage for *E. rudis* and *E. camaldulensis,* respectively). Based on the *k-*mer profiling analyses, the haplotype genome size of *E. rudis* was estimated to be 506 Mb while *E. camaldulensis* was estimated at a similar size of 510 Mb with a repeat content for both at around 36-37%. Heterozygosity was relatively high in both species (*E. camaldulensis*: 2.19%, *E. rudis*: 1.57%) and was expected to affect genome assembly [37] and estimated genome size [38].

#### Genome assemblies

We assembled the two eucalypts genomes with two assembly tools optimized for our data types: Canu [39] for CLR data and hifiasm [37] for HiFi data. The genome assembly statistics are shown in Table 1. The assembled diploid genome size based on the HiFi and on the CLR data is similar for *E. rudis* (975.2 Mb and 918.4 Mb, respectively) and *E. camaldulensis* (1053.1 Mb and 1093.7 Mb, respectively). The assembled *E. rudis* genome size is slightly lower than the *k*-mer profiling analysis which estimated a diploid size of slightly over 1012 Mb. For all assembly metrics, the HiFi assembly is superior to the CLR assembly for both *Eucalyptus* species, which is in agreement with HiFi-based assemblies for other species [8,39,40]. The contig L50 and N50 show noticeably higher contiguity in the HiFi assembly compared to the CLR assembly and the same trend is shown by the contig L90 and N90 statistics. For *E. rudis,* the number of contigs is approximately 75% higher in the CLR assembly than the HiFi assembly. This has a clear impact on the average contig length which is half that of the HiFi-based assembly. As for *E. camaldulensis*, the number of contigs and the average length were comparable between the HiFi and CLR assemblies. In addition, the longest contig (*E. rudis*: 61.8 Mb; *E. camaldulensis*: 69.1 Mb) was found in the HiFi assembly, which is nearly double the longest contig identified in the CLR assembly (33.8 Mb and 58.1 Mb, respectively).

**Table 1:** Genome assembly statistics for two *Eucalyptus* species. N50 and N90 represent the length of the shortest contig at 50% or 90% of the total assembly size. L50 and L90 represent the number of contigs that add up to 50% or 90% of the assembled genome size. Both N and L metrics add/count top contigs that are ordered in a descending manner based on length. We calculated the assembly stats using quast. CLR-based assemblies were 3-cycle polished as detailed in the Methods.

| Plants | Library | Total assembly (Mb) | Number of contigs | N50 (Mb) | L50 | N90 (Mb) | L90 | Longest Contig (Mb) | Average Length (Mb) |
| --- | --- | --- | --- | --- | --- | --- | --- | --- | --- |
| *E. rudis* | HiFi<sup>1</sup> | 975.2 | 1005 | 10.7 | 14 | 1.15 | 129 | 61.8 | 0.97 |
|  | CLR<sup>2</sup> | 918.4 | 1760 | 5.9 | 28 | 0.18 | 749 | 33.8 | 0.52 |
| *E. camaldulensis* | HiFi<sup>1</sup> | 1053.1 | 1131 | 18.9 | 14 | 1.61 | 114 | 69.1 | 0.93 |
|  | CLR<sup>3</sup> | 1093.7 | 1348 | 8.1 | 26 | 0.55 | 233 | 58.1 | 0.81 |
| <sup>1</sup> HiFi data were assembled using hifiasm. <sup>2</sup> CLR coverage is 31x. <sup>3</sup> CLR coverage was 118x (readjust for correct genome size) |  |  |  |  |  |  |  |  |  |

The genome assembly haploid sizes were estimated by separating haplotypes and purging haplotigs, and the completeness was assessed using BUSCO scores (Supp. Tables 5 and 6). Based on the primary haplo-purged assembly, the haploid genome size of *E. rudis* varies between 518 Mb (CLR) and 549 Mb (HiFi) producing an estimated ploidy of 1.8N. The high complete BUSCO scores (> 96%) for the primary contigs and a relatively high score for the alternative contigs (CLR: 73.4%; HiFi: 87.2%) indicate a reasonable separation of the haplotypes. In addition, the alternative contig set for the CLR assembly shows a lower degree of completeness which further demonstrates the superiority of the HiFi assembly. The haploid genome size for *E. camaldulensis* is about 523 and 532 Mb based respectively on CLR and HiFi primary assemblies. Both the primary and the alternative contig sets show high BUSCO scores (primary: > 97%; alternative: > 93%) in both CLR and HiFi assemblies (Supp. Table 5). The estimated ploidy of the assembly after haplo-purging is over 1.9N which, taken together with the BUSCO scores, suggests that the assembly is comprehensive. Alignment to the *E. grandis* genome [29] shows that our HiFi *E. camaldulensis* assembly has 9 full chromosomes with the remaining two chromosomes spanned almost fully by two contigs each (Fig. 2). Similarly, the *E. rudis* HiFi assembly has five full chromosomes and the remaining chromosomes spanned almost fully by two to three contigs each (Fig. 2).

**Figure 2:**
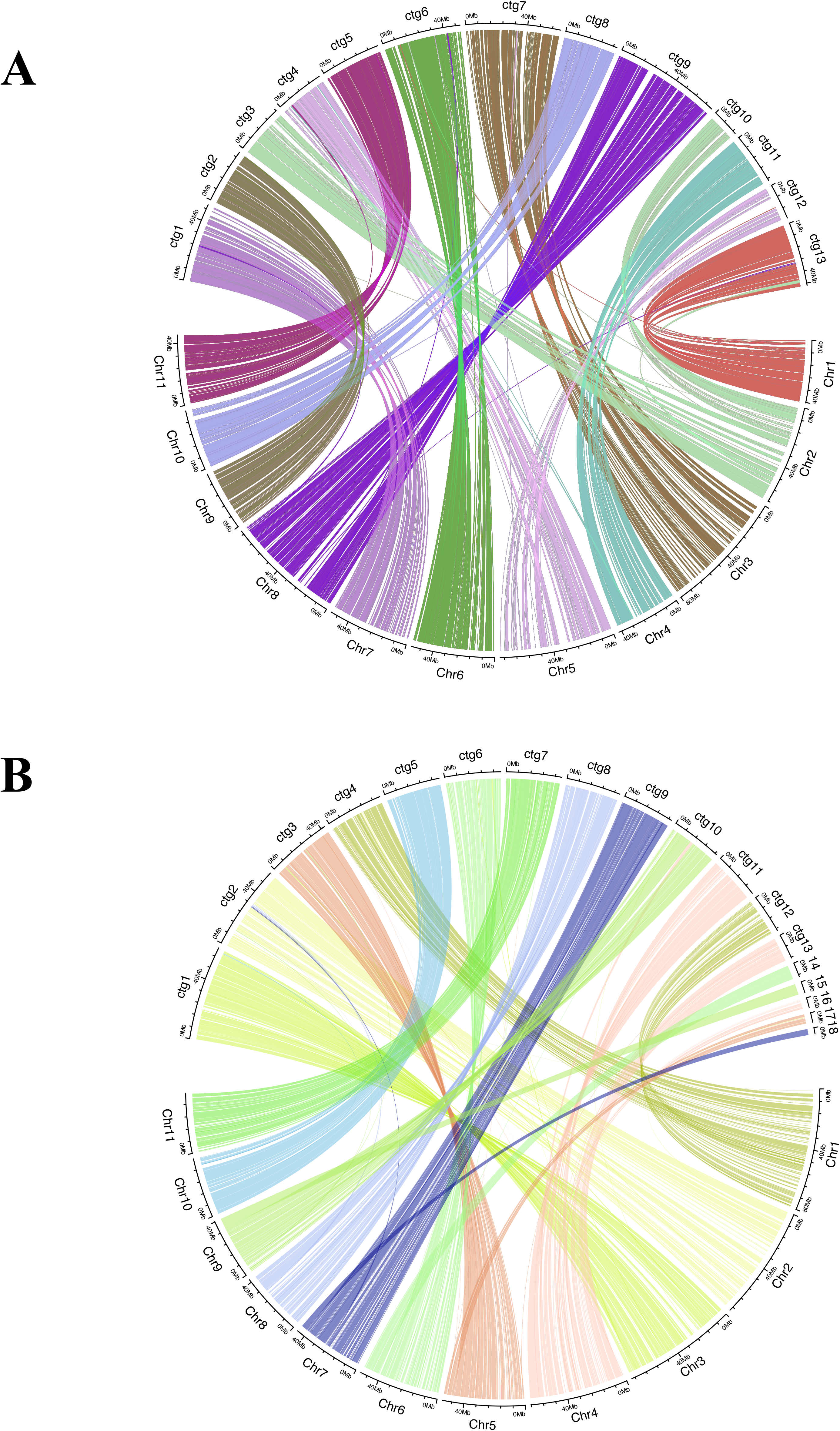
Alignment of *E. camaldulensis* (A) and *E. rudis* HiFi assemblies against *E. grandis* reference genome.

#### Genome assembly: computational resources

One of the main advantages for our chosen genome assembly workflow, using hifiasm with HiFi reads, are the savings in time and compute requirements, all with minimal manual intervention. The CLR data took significantly longer to assemble compared to the HiFi-based assemblies. At an estimated 27x CLR sequencing coverage, Canu took about 20 hours to finish in Cluster mode with 3444 CPU hours and a maximum RSS memory request peaking at 87GB. At a 127x CLR coverage, Canu, again in cluster mode, took 548 hours (1116 hours with queue wait and debugging) with over 72,491 CPU hours and maximum RSS request peaking at 69GB. While assembling the HiFi data using hifiasm took 80 mins for *E. rudis* (23x coverage) and 120mins for the *E. camaldulensis* (27x coverage). We ran hifiasm on a single 40 core node where it consumed 53 and 81 CPU hours for *E. rudis* and *E. camaldulensis*, respectively. In terms of memory, the former run peaked at 52 Gb and the latter at 64 Gb and but we observed memory overhead which therefore is a prerequisite for hifiasm. The generation of HiFi reads for *E. rudis* required 4.5 hours in cluster mode consuming 1,404 CPU hours over 3,212 total cores. For *E. camaldulensis*, HiFi generation needed 5 hours over 3,212 total cores using 2,000 CPU hours. The estimated total time from raw reads to HiFi data to the assembly of a high-quality contiguous draft for a haploid genome of 0.6 to 1.0 Gb is approximately one day. When combined with time estimates of HMW DNA extraction (one day), HiFi library preparation and sequencing (five days) and assembly; a high-quality draft genome can be prepared from plant samples in seven days, depending on available compute resources.

## Discussion

The current era of genomics is very promising with ambitious projects to sequence much of life on Earth [18,19,38]. The ideal goal for these projects would be to produce high-quality error-free, gapless or near-gapless haplotype-resolved genome assemblies. Very few gapless chromosome-level reference genomes today exist and typically require two or more different technologies like PacBio/ONT long reads, Illumina short and 10X linked reads sequencing, optical mapping, and Hi-C [5,9,11,41]. In particular, plants prove to have some of the most challenging genomes to assemble as they are rich in repetitive content, with high levels of heterozygosity and complex polyploidy [42]. Assembling a high quality plant genome with complete or near complete chromosomes therefore, requires high coverage, long read length and high quality sequences with a low error rate. Recently, PacBio has improved the circular consensus sequencing approach to generate long (≥ 15 Kb) HiFi reads with base-accuracy upwards of 99.8% [8], opening up the possibility of using only one technology to assemble a complete genome. We took advantage of this progress and developed a streamlined workflow able to produce a high-quality draft plant genome from plant tissue within seven days.

Our workflow is designed to be used as a rapid one-pass approach for assembling a high-quality draft genome that generates data suitable for most plant species. The use of one sequencing technology is a distinctive advantage of our workflow, although compatibility with long read ONT sequencing was demonstrated. The adoption of other sequencing technologies might improve the contiguity of a genome assembly, but a significant investment of additional resources may be needed, such as ultra-high molecular weight DNA extractions, increased wet-lab labor and time, additional equipment and reagents, various software packages and complex pipelines, and extensive bioinformatics analysis with manual intervention [5,9,11,38,41].

Despite the fact that our extraction approach relies on a column-based method and is thus susceptible to DNA shearing, we were able to deliver excellent quality DNA of high molecular weight. All the extracted samples had an average DNA length of over 80 Kb with optical density ratios indicating an absence of phenolics, carbohydrates, and protein and RNA contamination. We then incorporated two different methods of PacBio library preparation and sequencing, HiFi and CLR, allowing us to benchmark effectiveness and utility of each in turn. The former proved to generate high base-level accuracy long reads, while the latter produced subreads of considerable length. The averaged CLR N50 subread length was greater than 36 Kb on both Sequel I Sequel II platforms, exceeding therefore, the N50 of many recent studies [11,38,40,43].

We selected the CLR and HiFi data for *E. rudis* and *E. camaldulensis* to assess the final assembly step of our workflow. We then chose the best-in-class assembly packages for HiFi and CLR data [37,39] for our comparison. Although the eucalypt genomes assembled with both CLR and HiFi data are very contiguous, the higher base-level accuracy given by HiFi improves the assembly considerably thus removing the need for polishing with short read sequencing. In addition, our workflow emphasizes several strengths of the HiFi technology. Compared to the CLR assemblies, HiFi assemblies demanded less computational requirements, had higher BUSCO scores, showed several fold improvement of contig N50/N90 and L50/L90, and generated more complete genome assemblies, as has been previously published [8,37,40]. In fact, our HiFi sequencing data assembled with hifiasm, produced near-chromosome level haploid draft genomes. Our genomes are therefore comparable to single technology assemblies based on high coverage CLR and HiFi assemblies of homozygous maize strains and, more relevantly, to heterozygous plant species based on HiFi reads [10,37]. For a plant with a one Gb genome, we estimate that within approximately seven days a high quality draft genome assembly can be produced from plant tissue for an estimated consumables and compute cost of US$3000-US$4000 [5].

The two genomes sequenced here will improve our genomic knowledge of eucalypts, which at the moment is relatively sparse, and will assist with conservation issues and commercial uses. There are more than 800 eucalypt species, but only three genomes have been published: *E. grandis* [29], *E. pauciflora* [31] and *E. camaldulensis* [44]. The high-quality draft genome of *E. camaldulensis* generated here improves upon the published and highly fragmented genome produced by short read sequencing (Hirakawa et al. 2011), and the unpublished assembly recently produced from ONT long reads [45]. Based on assembly metrics (contig N50 of 18.9 Mb compared to 2.5 Mb for the ONT assembly) and our alignment to the *E. grandis* genome, indeed, suggest that our HiFi *E. camaldulensis* genome is a more contiguous assembly.

We also generated the first draft genome for *E. rudis*, a close relative of *E. camaldulensis* that inhabits a different ecological niche [46]. Our assembly of *E. rudis* produced with 23x HiFi coverage suggests a genome size of 549 Mb, similar to other eucalypts [29,31,44,45]. Moreover, the high BUSCO scores (> 96%), high quality assembly metrics and mostly complete chromosomes emphasize the high contiguity and quality of our draft genome.

The global initiatives to sequence and assemble genomes for many thousands of eukaryotic life forms, including plants, do not yet have a published standardized workflow [18,19,38,47,48]. Pursuing this goal, our workflow is an extremely valuable tool as it provides the foundation to produce high quality genome assemblies within seven days. Furthermore, this timeline might be further shortened by future bioinformatics and computational developments. Our workflow - and specifically our benchmarking efforts for DNA extraction, library preparation and sequencing, and genome assembly - will be a valuable resource for these consortia in the development of a fast and cost effective genome assembly pipeline. We envisage that our simplified and efficient workflow will be useful for plant researchers as well as specialized genome sequencing centers.

## Methods

### Sample Collection & HMW DNA Extraction

We chose seven plants from a broad range of taxa that are economically relevant, and represent typical species that may require high quality draft genomes (Supp. Table 1). Leaves were collected after at least 48 hours of dark treatment. To dark treat samples the entire plant, or a branch for large trees, was covered with light-opaque black plastic sheets with a few holes that allowed air flow. The leaves were sprayed with ethanol and wiped to remove contaminating organisms. Leaves were removed, and weighed before flash-freezing in liquid nitrogen.

HMW DNA was extracted by modifying a Qiagen Genomic protocol (Qiagen, Hilden, Germany). Briefly, frozen leaves were ground to a fine powder in a mortar and pestle under liquid nitrogen and stored at −80°C. For extraction, 1 g of ground leaf powder was resuspended in lysis buffer with 1 mg/ml of proteinase K (19133, Qiagen) and 190 μg/ml of RNase A (19101, Qiagen). The lysis solution was incubated at 50°C for at least 3.5 hours with gentle rocking. Following centrifugation for 15 min at room temperature at 3220 xg the supernatant was purified with genomic tip columns according to the manufacturer’s recommendations. After elution from the column the DNA was precipitated with 0.7 volumes of isopropanol and inverted slowly until the appearance of a floating DNA mass, or “jellyfish”. The DNA was hooked out with a plastic loop and washed in fresh 80% ethanol for 1 min three times. Repeated washing in ethanol has been shown to improve DNA purity without a decrease in yield [13]. The DNA pellet was resuspended in the EB buffer overnight at room temperature. The yield and quality of the DNA was assessed with a Broad Range dsDNA Qubit assay (Q32853, ThermoFisher, Waltham, MA, USA) and Nanodrop 8000 (ThermoFisher). In the event that absorbance ratios were not optimal [9,49], potentially indicating the presence of contaminants, a repeat bead clean was performed. To visualise the DNA smear pulse field gel electrophoresis (PFGE) was run with a 1% agarose TBE gel over 24 hours (Initial switch 1 sec, final switch 25 sec, 6 volt/cm, 120° included angle, Chef III, Bio-Rad, Hercules, CA, USA) with Midrange PFG and Lambda PFG markers (N0342S, N0341S, NEB, Ipswich, MA, USA). Alternatively, HMW DNA was also imaged with a gDNA 165 kb kit (FP-1002-0275) on the Femto Pulse system (Agilent, Santa Clara, CA, USA). A detailed procedure is deposited in protocols.io (dx.doi.org/10.17504/protocols.io.bafmibk6).

### PacBio and Oxford Nanopore Sequencing

#### CLR Library Preparation Sequencing with PacBio Sequel I and II

SMRTbell libraries were constructed with SMRTbell Express Template Prep Kit 2.0 (Pacific Biosciences, Menlo Park, CA, USA, 100-938-900). The input DNA, with a size distribution median predominantly above 80 Kb, was processed for SMRTbell library construction without any shearing (except for one library, Supp. Table 2). Initially, 15-20 μg of gDNA was cleaned-up and concentrated with 0.45x AMPure PB (Pacific Biosciences, 100-265-900), then 10 μg of gDNA was used in the first enzymatic reactions to remove single-strand overhangs followed by the DNA damage repair, end-repair/A-tailing reaction and finally, adapter ligation. The SMRTbell libraries were then purified with 0.45x AMPure PB beads before size selection on the BluePippin system (Sage Science, Beverley, MA, USA) with a 30 Kb cut-off using a 0.75% agarose cassette with U1 ladder. Following size selection, the libraries were given a final 1x AMPure PB bead clean-up and eluted in 10 μl of EB (Pacific Bioscience, 101-633-500). The concentration and size of SMRTbell were assessed with the Qubit dsDNA assay kit (Thermo Fisher Scientific, Waltham, MA, USA; Q32854), and the Genomic DNA 165 Kb Kit (Agilent; FP-1002-0275) on the FEMTO Pulse, respectively. For Sequel I, SMRTbell libraries were prepared for sequencing by annealing to Sequencing Primer v4 and Polymerase 3.0 with Sequel Sequencing kit 3.0 (Pacific Biosciences, 101-597-900) and Sequel Binding and Internal Control Kit 3.0 (Pacific Biosciences, 101-626-600); and SMRT Cell 1M v3 LR (Pacific Biosciences, 101-531-001) according to SMRT link Sample Setup v.7.0 instructions and sequencing for 10 or 20 hours without any pre-extension time. For Sequel II, SMRTbells were annealed with primer v4 and polymerase with Sequel II Binding Kit 2.0 and Internal Control Kit 1.0 (Pacific Biosciences,101-842-900), SMRT Cell 8M (Pacific Biosciences, 101-389-001) and according to SMRT link Sample Setup v.8.0 and sequence for 15 or 30 hours without any pre-extension time.

#### HiFi Library Preparation and Sequencing with PacBio Sequel II

15-20 μg of gDNA was diluted in 300 μl with 1x EB and was sheared using g-TUBE (Covaris, 520079) at 3500 - 4600 rpm in the Eppendorf 5424 (Eppendorf, Hamburg, Germany) for 2 minutes each spin. A repeat spin was implemented to make sure the entire gDNA had passed through the orifice. For shearing, small scale test shears were performed to make sure that the mode of the fragments were in the 15-20 Kb size range (size checked with FEMTO Pulse). A minimum 10 μg of sheared and 0.45x AMPure purified gDNA was carried into SMRTbell construction by using Express Template Prep Kit 2.0 + Enzyme Clean Up (101-843-100). Additional step of nuclease treatment of HiFi library after the ligation step was done to remove any non-intact SMRTbell templates. Following nuclease treatment, the SMRTbell was purified with two AMPure PB clean up steps, first with 0.45x, followed by 3.1x of diluted 35% v/v AMPure PB beads. The concentration and size of HiFi SMRTbell were assessed with Qubit dsDNA assay kit and gDNA 165kb kit of FEMTO Pulse respectively. The SMRTbell libraries were annealed and bound with sequencing primer v2 (101-847-900), and Sequel II DNA polymerase 2.0 from Sequel II Binding kit 2.0, 101-842-900, respectively using conditions specified in SMRT Link Sample Setup v.8.0. The final sample bound complex was sequenced with Sequel II Sequencing Kit 2.0 (101-820-200), and SMRT cell 8M Tray (101-389-001), and ran for 30 hours with 2 or 4 hours of pre-extension.

#### Oxford Nanopore Sequencing

Oxford Nanopore sequencing was performed on the GridION sequencer only for the two *Eucalyptus* species (*E. camaldulensis* and *E. rudis*). Prior to library preparation short fragments were depleted from the extracted HMW DNA using BluePippin with a 30 Kb cut-off or the Short Read Eliminator XL kit (SRE XL, Circulomics, Baltimore, MD, USA) using the manufacturer’s instructions. The size-selected DNA was cleaned-up with AMPure XP beads, and then processed into libraries using genomic DNA by ligation protocol (SQK-LSK109, Oxford Nanopore, Oxford, UK). Briefly, DNA (*E. camaldulensis*: 1.3 μg; *E. rudis*: 4 μg) was repaired and end-prepped prior to bead clean-up and adapter ligation. The ligated product is bead cleaned with Long Fragment Buffer (LFB) and eluted. The prepared library (*E. camaldulensis*: 8 fmol; *E. rudis*: 50 fmol) is then loaded onto GridION (FLO-MIN106D).

### Bioinformatics and Genome Assembly

#### Sequencing Data Analysis

For PacBio SMRTLink v8.0 was used for designing and monitoring sequencing runs, and analyzing and managing sequence data. For ONT MinKNOW Core 3.6.0 was used for data acquisition and real-time analysis. Reads were base-called using Guppy 3.2.8 from FAST5 files to produce FASTQ files. Statistical analyses were performed using R 3.6.1 [50].

The quality metrics of basecalled reads were also calculated using LongQC version 1.2 [35]. We applied pb-hifi, pb-sequel, and ont-ligation profiles for PacBio HiFi, PacBio CLR, and Oxford Nanopore datasets, respectively.

#### Genome Size Estimation

GenomeScope 2.0 [36] was used to estimate, *in silico*, the genome sizes of both *Eucalyptus* species. The software was run with the following parameters: [*k-mer length*=*12, Ploidy*=*2, Max k-mer coverage*=−*1, average k-mer coverage for polyploid genome*=−*1*]. We calculated the k-mer distribution, which we then fed to GenomeScope, using JellyFish [51] with the following parameters: [*jellyfish count* -*C* -*m 21* -*s 1000000000*].

#### Genome assemblies

We assembled the HiFi data for both *Eucalyptus* species using the newly-released assembler hifiasm v0.8 - r279 [37]. We run hifiasm with default settings (*-r3 -a4 -k51 -w51 -f37 -D5.0 -N100 -z0 -m10000000 -p 100000 -n3 -x0.8 -y0.2*). We also separated haplotigs using the purge-dups module in hifiasm using default settings (*-l2 -s0.75 -O1*). Output from hifiasm is two GFA graph files: one for the primary contigs and another for the alternative haplotigs. We converted the GFA files to the FASTA format using gfatools v0.4-r179-dirty [52]. To produce the total assembly stats, we remerged the primary and alternative haplotigs using seqkit [53].

For PacBio CLR we used the Canu assembler v2 (github r9818) [39]. To force Canu to keep haplotigs separate, we set the following parameters: *batOptions*=-*dg 3 -db 3 -dr 1 -ca 500 -cp 50*. For speed gains, we run Canu in cluster mode. Where deep coverage was available, we increased the amount of data used in the assembly beyond the default and commonly used 40x coverage.

#### Polishing of genome assemblies

We polished the CLR-based assemblies using the Arrow algorithm in the gcpp tool from PacBio’s SMRT Link v8.0 stack. First, we aligned the raw CLR data against the initial assembly using pbmm2 v1.1.0, which is a version of Minimap2 [54] adapted to PacBio’s native format. The alignment is then used for consensus calling and polishing using gcpp v1.0.0. We repeated the process for two additional polishing cycles whereby we feed the polished assembly from the previous cycle as the alignment reference in the next cycle. The HiFi-based assemblies do not require additional polishing to the highly-accurate starting material.

#### Genome assemblies’ assessment

We generated comprehensive assembly statistics using QUAST-LG v5.0.2 [55]. To assess the biological integrity of the assemblies, we used BUSCO v3 [56] as a proxy of genome completeness.

#### Haplotig purging

We removed haplotigs for the CLR-based assemblies using purge_dups [57] using default settings. We manually inspected read depth to adjust coverage cut-offs where necessary for best performance. The hifiasm pipeline, on the other hand, is able to separate haplotigs for the HiFi-based assemblies without additional tools. We validated the success of this step for all assemblies using BUSCO v3 [56].

We estimated the ploidy (EP) of the assembly using the following basic formulae:

*EP = Smaller haploid / Bigger haploid*. The smaller and bigger haploids refer to either the primary or the alternative set of contigs from the haplotigs purging step. This remains an estimate and should be taken as such; because we naively assume that the larger set of haplotigs constitutes a complete haploid N.

#### Phenotypic and *in silico* identification of the *Eucalyptus* species

Phenotypic identification of the two *Eucalyptus* species indicated the two sequenced trees belonged to *Eucalyptus camaldulensis* subspecies *obtusa* and *Eucalyptus rudis* subspecies *rudis*. The species were identified by examination of leaves, flower buds, seed pods, seeds, bark and tree morphology. The phenotypic identification was then confirmed to species level by *in silico* DNA analysis. *in silico* DNA metabarcoding based on 336 complete ITS sequences from different *Eucalyptus* genera indicated that the best hits of our query sequences were *Eucalyptus camaldulensis* and *Eucalyptus rudis* for each dataset (Supp. Fig. 12).

We obtained 335 complete ITS sequences representing 304 different *Eucalyptus* genera and 31 unknown from the PLANiTS dataset version 29-03-2020 [58].

As the dataset did not include the ITS sequence for *E. rudis* we downloaded it separately from NCBI Nucleotide (accession: KT631323.1) bringing the total number of our *Eucalyptus* ITS sequences to 336. We also replaced the ITS sequence for *E. camaldulensis* with the longer version deposited in the NCBI nucleotide database (accession: AF190363.1) which, like the ones for *E. rudis*, also includes partial 18S, full 5.8S and partial 26S rRNA genes.

We aligned both *Eucalyptus* genomes against the ITS dataset using blastn v2.7.1 [59] with the E-value cutoff set to 1e-5. We assessed the top hits taking into account percentage identity, length of alignment, mismatches and gaps.

### Data availability

The assembled genomes and relevant raw data are available under NCBI Project ID PRJNA674723.

## Acknowledgements

We thank Dr. Boubacar Kountche and Prof. Salim AlBabili for insightful discussions on the wet-lab work and their constructive support for BCL, and the KAUST Supercomputing Core Lab for providing computing resources and advice. We thank Dr. Dean Nicolle for the phenotypic identification of the two *Eucalyptus* species. This study used resources provided by the KAUST Core Labs and we thank the Core Labs management for supporting this activity. In fond memory of our talented colleague Dr Kamel Jabbari.

## Author’s Contributions

P.D, K.C., and A.P performed laboratory work, PacBio and ONT sequencing, and interpreted the results. S.B. carried out the genome assemblies and all bioinformatics analyses with support from Y.F.. K.J. estimated size and the heterozygosity level on the unassembled HiFi sequencing data. M.P. and R.S. provided plant samples for sequencing. N.C. supported the study and provided managerial guidance. Y.F. performed LongQC analyses, supervised bioinformatic analysis and provided valuable advice. L.E. conceived and planned this study, interpreted the results, and supervised all aspects of this work. L.E., P.D. and S.B. wrote the manuscript with input from all authors. All authors read, edited, and approved the final manuscript for submission.

## Conflict of Interest

The authors declare that they have no conflict of interest.

## Notes

### Competing Interest Statement

The authors have declared no competing interest.

## References

1. Burgess MG, Gaines SD. The scale of life and its lessons for humanity. Proc Natl Acad Sci. 2018;115:6328–30.

2. Shukla PR, Skea J, Buendia EC, Masson-Delmotte V, Pörtner H-O, Roberts DC, et al. IPCC, 2019: Climate Change and Land: an IPCC special report on climate change, desertification, land degradation, sustainable land management, food security, and greenhouse gas fluxes in terrestrial ecosystems. In press.

3. Sala OE, Chapin FS, Iii, Armesto JJ, Berlow E, Bloomfield J, et al. Global Biodiversity Scenarios for the Year 2100. Science. 2000;287:1770–4.

4. Surfing the genomic new wave. Nat Plants. 2018;4:393–393.

5. Logsdon GA, Vollger MR, Eichler EE. Long-read human genome sequencing and its applications. Nat Rev Genet [Internet]. 2020 [cited 2020 Jun 7]; Available from: http://www.nature.com/articles/s41576-020-0236-x

6. Miga KH, Koren S, Rhie A, Vollger MR, Gershman A, Bzikadze A, et al. Telomere-to-telomere assembly of a complete human X chromosome. bioRxiv. 2019;735928.

7. Jain M, Koren S, Miga KH, Quick J, Rand AC, Sasani TA, et al. Nanopore sequencing and assembly of a human genome with ultra-long reads. Nat Biotechnol. 2018;36:338–45.

8. Wenger AM, Peluso P, Rowell WJ, Chang P-C, Hall RJ, Concepcion GT, et al. Accurate circular consensus long-read sequencing improves variant detection and assembly of a human genome. Nat Biotechnol. 2019;37:1155–62.

9. Jung H, Winefield C, Bombarely A, Prentis P, Waterhouse P. Tools and Strategies for Long-Read Sequencing and De Novo Assembly of Plant Genomes. Trends Plant Sci. 2019;24:700–24.

10. Ou S, Liu J, Chougule KM, Fungtammasan A, Seetharam AS, Stein JC, et al. Effect of sequence depth and length in long-read assembly of the maize inbred NC358. Nat Commun. Nature Publishing Group; 2020;11:2288.

11. Liu J, Seetharam AS, Chougule K, Ou S, Swentowsky KW, Gent JI, et al. Gapless assembly of maize chromosomes using long-read technologies. Genome Biol. 2020;21:121.

12. Inglis PW, Pappas M de CR, Resende LV, Grattapaglia D. Fast and inexpensive protocols for consistent extraction of high quality DNA and RNA from challenging plant and fungal samples for high-throughput SNP genotyping and sequencing applications. Kalendar R, editor. PLOS ONE. 2018;13:e0206085.

13. Mayjonade B, Gouzy J, Donnadieu C, Pouilly N, Marande W, Callot C, et al. Extraction of high-molecular-weight genomic DNA for long-read sequencing of single molecules. BioTechniques [Internet]. 2016 [cited 2019 May 27];61. Available from: https://www.future-science.com/doi/10.2144/000114460

14. Vaillancourt B, Buell CR. High molecular weight DNA isolation method from diverse plant species for use with Oxford Nanopore sequencing. bioRxiv. 2019;783159.

15. Varma A, Padh H, Shrivastava N. Plant genomic DNA isolation: An art or a science. Biotechnol J. 2007;2:386–92.

16. Zhang M, Zhang Y, Scheuring CF, Wu C-C, Dong JJ, Zhang H-B. Preparation of megabase-sized DNA from a variety of organisms using the nuclei method for advanced genomics research. Nat Protoc. 2012;7:467–78.

17. Healey A, Furtado A, Cooper T, Henry RJ. Protocol: a simple method for extracting next-generation sequencing quality genomic DNA from recalcitrant plant species. Plant Methods. 2014;10:21.

18. Lewin HA, Robinson GE, Kress WJ, Baker WJ, Coddington J, Crandall KA, et al. Earth BioGenome Project: Sequencing life for the future of life. Proc Natl Acad Sci. 2018;115:4325–33.

19. Cheng S, Melkonian M, Smith SA, Brockington S, Archibald JM, Delaux P-M, et al. 10KP: A phylodiverse genome sequencing plan. GigaScience [Internet]. Oxford Academic; 2018 [cited 2020 Nov 26];7. Available from: https://academic.oup.com/gigascience/article/7/3/giy013/4880447

20. Debieu M, Kanfany G, Laplaze L. Pearl Millet Genome: Lessons from a Tough Crop. Trends Plant Sci. 2017;22:911–3.

21. Jiao Y, Peluso P, Shi J, Liang T, Stitzer MC, Wang B, et al. Improved maize reference genome with single-molecule technologies. Nature. 2017;546:524–7.

22. Liu J, Seetharam AS, Chougule K, Ou S, Swentowsky KW, Gent JI, et al. Gapless assembly of maize chromosomes using long read technologies [Internet]. Bioinformatics; 2020 Jan. Available from: http://biorxiv.org/lookup/doi/10.1101/2020.01.14.906230

23. Schnable PS, Ware D, Fulton RS, Stein JC, Wei F, Pasternak S, et al. The B73 maize genome: complexity, diversity, and dynamics. Science. 2009;326:1112–5.

24. Varshney RK, Shi C, Thudi M, Mariac C, Wallace J, Qi P, et al. Pearl millet genome sequence provides a resource to improve agronomic traits in arid environments. Nat Biotechnol. 2017;35:969–76.

25. Pearlstein SL, Felger RS, Glenn EP, Harrington J, Al-Ghanem KA, Nelson SG. Nipa (*Distichlis palmeri*): A perennial grain crop for saltwater irrigation. J Arid Environ. 2012;82:60–70.

26. Glenn EP, Anday T, Chaturvedi R, Martinez-Garcia R, Pearlstein S, Soliz D, et al. Three halophytes for saline-water agriculture: An oilseed, a forage and a grain crop. Environ Exp Bot. 2013;92:110–21.

27. Reddy MP, Shah MT, Patolia JS. Salvadora persica, a potential species for industrial oil production in semiarid saline and alkali soils. Ind Crops Prod. 2008;28:273–8.

28. Monfared MA, Samsampour D, Sharifi-Sirchi GR, Sadeghi F. Assessment of genetic diversity in *Salvadora persica L.* based on inter simple sequence repeat (ISSR) genetic marker. J Genet Eng Biotechnol. 2018;16:661–7.

29. Myburg AA, Grattapaglia D, Tuskan GA, Hellsten U, Hayes RD, Grimwood J, et al. The genome of *Eucalyptus grandis*. Nature. 2014;510:356–62.

30. Schalamun M, Nagar R, Kainer D, Beavan E, Eccles D, Rathjen JP, et al. Harnessing the MinION: An example of how to establish long-read sequencing in a laboratory using challenging plant tissue from *Eucalyptus pauciflora*. Mol Ecol Resour. 2019;19:77–89.

31. Wang W, Das A, Kainer D, Schalamun M, Morales-Suarez A, Schwessinger B, et al. The draft nuclear genome assembly of *Eucalyptus pauciflora*: a pipeline for comparing de novo assemblies. GigaScience [Internet]. 2020 [cited 2020 Jan 12];9. Available from: https://academic.oup.com/gigascience/article/9/1/giz160/5694103

32. Simbolo M, Gottardi M, Corbo V, Fassan M, Mafficini A, Malpeli G, et al. DNA Qualification Workflow for Next Generation Sequencing of Histopathological Samples. PLOS ONE. 2013;8:e62692.

33. Guide - Step-By-Step Run Performance Evaluation. 2020;15.

34. Belser C, Istace B, Denis E, Dubarry M, Baurens F-C, Falentin C, et al. Chromosome-scale assemblies of plant genomes using nanopore long reads and optical maps. Nat Plants. Nature Publishing Group; 2018;4:879–87.

35. Fukasawa Y, Ermini L, Wang H, Carty K, Cheung M-S. LongQC: A Quality Control Tool for Third Generation Sequencing Long Read Data. G3amp58 GenesGenomesGenetics. 2020;10:1193–6.

36. Ranallo-Benavidez TR, Jaron KS, Schatz MC. GenomeScope 2.0 and Smudgeplot for reference-free profiling of polyploid genomes. Nat Commun. Nature Publishing Group; 2020;11:1432.

37. Cheng H, Concepcion GT, Feng X, Zhang H, Li H. Haplotype-resolved de novo assembly with phased assembly graphs. ArXiv200801237 Q-Bio [Internet]. 2020 [cited 2020 Aug 26]; Available from: http://arxiv.org/abs/2008.01237

38. Rhie A, McCarthy SA, Fedrigo O, Damas J, Formenti G, Koren S, et al. Towards complete and error-free genome assemblies of all vertebrate species. bioRxiv. Cold Spring Harbor Laboratory; 2020;2020.05.22.110833.

39. Nurk S, Walenz BP, Rhie A, Vollger MR, Logsdon GA, Grothe R, et al. HiCanu: accurate assembly of segmental duplications, satellites, and allelic variants from high-fidelity long reads. Genome Res [Internet]. 2020 [cited 2020 Sep 9]; Available from: http://genome.cshlp.org/content/early/2020/09/02/gr.263566.120

40. Vollger MR, Logsdon GA, Audano PA, Sulovari A, Porubsky D, Peluso P, et al. Improved assembly and variant detection of a haploid human genome using single-molecule, high-fidelity long reads. Ann Hum Genet. 2020;84:125–40.

41. Logsdon GA, Vollger MR, Hsieh P, Mao Y, Liskovykh MA, Koren S, et al. The structure, function, and evolution of a complete human chromosome 8. bioRxiv. Cold Spring Harbor Laboratory; 2020;2020.09.08.285395.

42. Mishra DC, Lal SB, Sharma A, Kumar S, Budhlakoti N, Rai A. Strategies and Tools for Sequencing and Assembly of Plant Genomes. In: Kumar Chakrabarti S, Xie C, Kumar Tiwari J, editors. Potato Genome [Internet]. Cham: Springer International Publishing; 2017 [cited 2020 Nov 1]. p. 81–93. Available from: https://doi.org/10.1007/978-3-319-66135-3_5

43. Zhou Y, Chebotarov D, Kudrna D, Llaca V, Lee S, Rajasekar S, et al. A platinum standard pan-genome resource that represents the population structure of Asian rice. Sci Data. Nature Publishing Group; 2020;7:113.

44. Hirakawa H, Nakamura Y, Kaneko T, Isobe S, Sakai H, Kato T, et al. Survey of the genetic information carried in the genome of *Eucalyptus camaldulensis*. Plant Biotechnol. 2011;28:471–80.

45. ASM1418270v1 - Genome - Assembly - NCBI [Internet]. [cited 2020 Nov 3]. Available from: https://www.ncbi.nlm.nih.gov/assembly/GCA_014182705.1

46. Boland DJ, Brooker MIH, Chippendale GM, Hall N, Hyland BPM, Johnston RD, et al. Forest Trees of Australia. Csiro Publishing; 2006.

47. Darwin Tree Of Life [Internet]. [cited 2020 Nov 18]. Available from: https://www.darwintreeoflife.org/

48. Eucalyptus (ID 509734) - BioProject - NCBI [Internet]. [cited 2020 May 17]. Available from: https://www.ncbi.nlm.nih.gov/bioproject/509734

49. Pacific Biosciences. Technical Note: Preparing DNA for PacBio HiFi- Sequencing Extraction and Quality Control [Internet]. Prep. DNA PacBio HiFi Seq. — Extr. Qual. Control. 2020 [cited 2020 Sep 30]. Available from: https://www.pacb.com/wp-content/uploads/Technical-Note-Preparing-DNA-for-PacBio-HiFi-Sequencing-Extraction-and-Quality-Control.pdf

50. R Core Team. R: A Language and Environment for Statistical Computing [Internet]. Vienna, Austria: R Foundation for Statistical Computing; 2017. Available from: https://www.R-project.org/

51. Marçais G, Kingsford C. A fast, lock-free approach for efficient parallel counting of occurrences of k-mers. Bioinformatics. Oxford Academic; 2011;27:764–70.

52. Li H, Feng X, Chu C. The design and construction of reference pangenome graphs with minigraph. Genome Biol. 2020;21:265.

53. Shen W, Le S, Li Y, Hu F. SeqKit: A Cross-Platform and Ultrafast Toolkit for FASTA/Q File Manipulation. PLOS ONE. Public Library of Science; 2016;11:e0163962.

54. Li H. Minimap2: pairwise alignment for nucleotide sequences. Bioinformatics. Oxford Academic; 2018;34:3094–100.

55. Mikheenko A, Prjibelski A, Saveliev V, Antipov D, Gurevich A. Versatile genome assembly evaluation with QUAST-LG. Bioinformatics. Oxford Academic; 2018;34:i142–50.

56. Seppey M, Manni M, Zdobnov EM. BUSCO: Assessing Genome Assembly and Annotation Completeness. In: Kollmar M, editor. Gene Predict [Internet]. New York, NY: Springer New York; 2019 [cited 2020 Aug 26]. p. 227–45. Available from: http://link.springer.com/10.1007/978-1-4939-9173-0_14

57. Guan D, McCarthy SA, Wood J, Howe K, Wang Y, Durbin R. Identifying and removing haplotypic duplication in primary genome assemblies. Bioinformatics. Oxford Academic; 2020;36:2896–8.

58. Banchi E, Ametrano CG, Greco S, Stanković D, Muggia L, Pallavicini A. PLANiTS: a curated sequence reference dataset for plant ITS DNA metabarcoding. Database [Internet]. 2020 [cited 2020 Dec 22];2020. Available from: https://doi.org/10.1093/database/baz155

59. Altschul SF, Gish W, Miller W, Myers EW, Lipman DJ. Basic local alignment search tool. J Mol Biol. England; 1990;215:403–10.

