## Supplementary Tables for "LeafGo: Leaf to Genome, a quick workflow to produce high-quality *De novo* genomes with Third Generation Sequencing technology"

#### Supplementary Table 1: General information of the seven study plants, and the yield and quality of extracted HMW DNA

Taxonomic details, and the estimated genome size of the plants used in this study as well as yield and quality of extracted HMW DNA determined from Qubit and Nanodrop analysis.

| **Species** | **Common Name** | **Family** | **Plant type** | **Genome Size (Mb) *estimation** | **DNA yield (average ± SD) N, min - max (µg)** | **$A_{260/280}$ Average ±SD** | **$A_{260/230}$ average±SD** |
|---|---|---|---|---|---|---|---|
| *Pennisetum glaucum* | Pearl Millet | Poaceae | Grass | 1816 | 31.3 ± 9.2<br>4, 18.3 - 38.8 | 1.83 ± 0.00 | 2.27 ± 0.15 |
| *Salvadora persica* | Toothbrush tree | Salvadoraceae | Small evergreen tree | 412 | 161.3 ± 27.3<br>2, 142.0-180.6 | 1.86 ± 0.01 | 2.20 ± 0.13 |
| *Eucalyptus rudis* | Flooded gum | Myrtaceae | Hardwood tree | 600* | 109.5 ± 104.8<br>6, 15.0 - 278.4 | 1.85 ± 0.04 | 2.30 ± 0.17 |
| *Eucalyptus camaldulensis* | River red gum | Myrtaceae | Hardwood tree | 600* | 70.1 ± 53.9<br>4, 30.8 - 150.0 | 1.84 ± 0.03 | 2.26 ± 0.09 |
| *Salicornia bigelovii* | Dwarf saltwort | Amaranthaceae | Annual shrub | 1300* | 21.0<br>1, NA | 1.78 | 2.10 |
| *Distichlis palmeri* | Nipa grass | Poaceae | Saltgrass | 400* | 118.3 ± 51.9<br>2, 81.6 - 155.0 | 1.83 ± 0.02 | 2.06 ± 0.05 |
| *Zea mays* | Sweetcorn | Poaceae | Grass | 2135 | 140<br>1, NA | 1.82 | 2.26 |

### Supplementary Table 2: PacBio sequencing results for seven different plant species

| Sample Details | Sequel Platform | Type | Insert Size (Kb) | Movie Time (hours) | Total Bases (Gb) | Q20 Yield (Gb) | Q20 Read Quality (median) | Polymerase av. (bp) | Polymerase N50 (bp) | Subread av. (bp) | Subread N50 (bp) | P0 % | P1 % | P2 % | Control Polymerase Read Length (Mean bp) | Local Base Rate |
|---|---|---|---|---|---|---|---|---|---|---|---|---|---|---|---|---|
| *D. palmeri* | I | CLR | 30 | 20 | 14.1 | NA | NA | 23094 | 39464 | 21004 | 35435 | 20 | 62 | 18 | 33896 | 2.80 |
| *D. palmeri* | I | CLR | 30 | 20 | 15.2 | NA | NA | 21973 | 37524 | 20255 | 34414 | 13 | 71 | 16 | 46266 | 2.79 |
| *E. camaldulensis* | I | CLR | 30 | 20 | 18.1 | NA | NA | 27292 | 47335 | 23299 | 38312 | 19 | 67 | 14 | 39479 | 2.80 |
| *E. camaldulensis* | I | CLR | 30 | 20 | 15.3 | NA | NA | 22041 | 37357 | 20477 | 34503 | 12 | 71 | 17 | 44527 | 2.73 |
| *E. rudis* | I | CLR | 30 | 20 | 17.8 | NA | NA | 27719 | 47854 | 24691 | 40998 | 21 | 64 | 14 | 37377 | 2.82 |
| *E. rudis* | I | CLR | 30 | 20 | 15.7 | NA | NA | 23465 | 41037 | 22080 | 38105 | 15 | 69 | 16 | 40673 | 2.63 |
| *P. glaucum* | I | CLR | 30 | 20 | 11.1 | NA | NA | 28600 | 47852 | 27046 | 44686 | 51 | 42 | 7 | 51922 | 2.57 |
| *P. glaucum* | I | CLR | 30 | 20 | 10.3 | NA | NA | 23480 | 41756 | 22570 | 39976 | 46 | 47 | 7 | 47919 | 2.56 |
| *P. glaucum* | I | CLR | 30 | 10 | 11.0 | NA | NA | 24392 | 41309 | 23748 | 39877 | 44 | 47 | 9 | 36426 | 2.68 |
| *P. glaucum* | I | CLR | 30 | 20 | 13.8 | NA | NA | 24430 | 40484 | 23388 | 38522 | 34 | 58 | 7 | 47332 | 2.41 |
| *P. glaucum* | I | CLR | 30 | 10 | 10.2 | NA | NA | 19362 | 34312 | 19063 | 33654 | 34 | 54 | 11 | 30748 | 2.43 |
| *P. glaucum* | I | CLR | 30 | 20 | 9.7 | NA | NA | 19121 | 29486 | 18512 | 28583 | 38 | 53 | 10 | 50240 | 2.62 |
| *P. glaucum* | I | CLR | 30 | 10 | 8.1 | NA | NA | 17372 | 29929 | 16978 | 28915 | 43 | 48 | 9 | 37545 | 2.61 |
| *P. glaucum* | I | CLR | 30 | 10 | 3.9 | NA | NA | 14747 | 21877 | 14524 | 21522 | 57 | 28 | 15 | 31185 | 2.64 |
| *P. glaucum* | I | CLR | 30 | 20 | 8.9 | NA | NA | 29054 | 45656 | 27336 | 42461 | 61 | 33 | 7 | 48680 | 2.49 |
| *P. glaucum* | I | CLR | 30 | 20 | 3.3 | NA | NA | 24691 | 44270 | 22557 | 40287 | 66 | 16 | 18 | 42502 | 2.44 |
| *P. glaucum* | I | CLR | 30 | 20 | 3.6 | NA | NA | 23435 | 42508 | 21491 | 38828 | 76 | 18 | 6 | 33458 | 2.74 |
| *P. glaucum** | I | CLR | 30 | 20 | 14.8 | NA | NA | 22964 | 38087 | 20031 | 33199 | 21 | 65 | 14 | 40927 | 2.77 |
| *S. bigelovii* | I | CLR | 30 | 20 | 17.0 | NA | NA | 31492 | 49431 | 26832 | 41558 | 32 | 56 | 12 | 45873 | 2.83 |
| *S. persica* | I | CLR | 30 | 20 | 4.3 | NA | NA | 28105 | 46406 | 26166 | 42980 | 75 | 18 | 6 | 34346 | 2.89 |
| *S. persica* | I | CLR | 30 | 20 | 14.2 | NA | NA | 20041 | 35828 | 18734 | 33322 | 12 | 73 | 15 | 45315 | 2.61 |
| *Z. mays* | I | CLR | 30 | 20 | 12.5 | NA | NA | 22897 | 38381 | 21333 | 35909 | 27 | 58 | 15 | 39190 | 2.63 |
| *E. camaldulensis* | II | CLR | 30 | 15 | 195.4 | NA | NA | 28350 | 45997 | 25489 | 39777 | 11 | 86 | 3 | 31888 | 2.01 |
| *E. rudis* | II | CLR | 30 | 30 | 150.6 | NA | NA | 27593 | 42816 | 22857 | 35313 | 30 | 68 | 2 | 53880 | 2.05 |
| *E. rudis* | II | CLR | 30 | 30 | 140.8 | NA | NA | 24789 | 36623 | 21494 | 32383 | 28 | 71 | 2 | 52398 | 2.07 |
| *S. persica* | II | CLR | 30 | 15 | 183.7 | NA | NA | 27773 | 44723 | 24753 | 38085 | 14 | 83 | 3 | 35769 | 2.11 |
| *D. palmeri* | II | HiFi | 18 | 30 | 113.9 | 7.6 | 33 | 107849 | 199058 | 17750 | 22156 | 86 | 13 | 0 | 61220 | 2.21 |
| *D. palmeri* | II | HiFi | 18 | 30 | 356.3 | 23.0 | 33 | 96533 | 189596 | 15300 | 19603 | 52 | 46 | 2 | 63774 | 2.25 |
| *E. camaldulensis* | II | HiFi | 17 | 30 | 477.3 | 28.7 | 33 | 88261 | 175407 | 14568 | 17673 | 29 | 68 | 3 | 56714 | 2.20 |
| *E. rudis* | II | HiFi | 17 | 30 | 317.2 | 22.1 | 32 | 101976 | 195495 | 17921 | 21714 | 60 | 39 | 1 | 55468 | 2.22 |
| *S. persica* | II | HiFi | 16 | 30 | 455.4 | 29.4 | 36 | 118199 | 200898 | 15723 | 17626 | 51 | 48 | 1 | 59746 | 2.25 |

* Representative sample of sheared gDNA for library preparation

**Supplementary Table 3: Correlation between library loading and throughput and N50**

Cells shaded in red represent significant results (p-value ≤ 0.05).

| Spearman's rho | P0% | P1% | P2% | Longest Subread N50 (bp) | Total Bases (Gb) | Polymerase N50 (bp) |
|---|---|---|---|---|---|---|
| **P0%** | 1 | -0.98 | -0.63 | 0.42 | -0.84 | 0.26 |
| **P1%** | -0.98 | 1 | 0.52 | -0.39 | 0.86 | -0.23 |
| **P2%** | -0.63 | 0.52 | 1 | -0.42 | 0.39 | -0.3 |
| **Longest Subread N50 (bp)** | 0.42 | -0.39 | -0.42 | 1 | 0 | 0.93 |
| **Total Bases (Gb)** | -0.84 | 0.86 | 0.39 | 0 | 1 | 0.22 |
| **Polymerase N50 (bp)** | 0.26 | -0.23 | -0.3 | 0.93 | 0.22 | 1 |
| **p-value** | **P0%** | **P1%** | **P2%** | **Longest Subread N50 (bp)** | **Total Bases (Gb)** | **Polymerase N50 (bp)** |
| **P0%** | | 0.0000 | 0.0018 | 0.0499 | 0.0000 | 0.2463 |
| **P1%** | 0.0000 | | 0.0129 | 0.0717 | 0.0000 | 0.306 |
| **P2%** | 0.0018 | 0.0129 | | 0.0548 | 0.0763 | 0.1752 |
| **Longest Subread N50 (bp)** | 0.0499 | 0.0717 | 0.0548 | | 0.9861 | 0.0000 |
| **Total Bases (Gb)** | 0.0000 | 0.0000 | 0.0763 | 0.9861 | | 0.3159 |
| **Polymerase N50 (bp)** | 0.2463 | 0.306 | 0.1752 | 0.0000 | 0.3159 | |

**Supplementary Table 4: Oxford Nanopore Technology Sequencing Results for two *Eucalyptus* species**

| Species | Active pores | Run time (hours) | Yield (Gb) | Normalized yield (Mb/ hour) | N50 (Kb) | Size selection method |
|---|---|---|---|---|---|---|
| *E. camaldulensis* | 1229 | 24 | 8.9 | 370.8 | 37.2 | SRE-XL |
| *E. camaldulensis* | 1095 | 24 | 6.2 | 258.3 | 43.8 | BluePippin 30Kb |
| *E. rudis* | 1679 | 48 | 13.7 | 285.4 | 42.8 | BluePippin 30Kb |
| *E. rudis* | 1699 | 48 | 16.2 | 337.5 | 41.8 | SRE-XL |

### Supplementary Table 5: Haplotype-separated assembly stats.

**P**: primary haplotigs/contigs which refer to the longer of the contigs that belong to a region of high heterozygosity. A: alternative haplotigs/contigs are the other non-primary haplotigs/contigs. **S**: single complete BUSCO, **D**: duplicate complete BUSCO.

| **Species** | **Assembly type** | **Primary/ Alternative** | **Total Length (Mb)** | **N50 (Mb)** | **Longest (Mb)** | **Ploidy (2N)[ß]** | **BUSCO** | | |
|---|---|---|---|---|---|---|---|---|---|
| | | | | | | | **Complete** | **Fragment** | **Missing** |
| *Eucalyptus rudis* | HiFi [∂] | P | 549 | 36.08 | 61.81 | 1.77N | 97.3 [S:91.7,D:5.6] | 1.1 | 1.6 |
| | | A | 425 | 3.5 | 10.74 | | 87.2 [S:83.4,D:3.8] | 1.3 | 11.5 |
| | CLR [Δ] | P | 518 | 16.34 | 33.77 | 1.77N | 96.3 [S:92.4,D:3.9] | 1.7 | 2 |
| | | A | 399 | 0.37 | 3.88 | | 73.4 [S:66.7,D:6.7] | 2.3 | 24.3 |
| *E. camaldulensis* | HiFi [ø] | P | 532 | 41.47 | 69.1 | 1.98N | 97.2 [S:93.9,D:3.3] | 0.9 | 1.9 |
| | | A | 520 | 4.1 | 19.36 | | 94.2 [S:89.7,D:4.5] | 1 | 4.8 |
| | CLR [Δ] | P | 523 | 29.31 | 58 | 1.92N | 97.3 [S:93.2,D:4.1] | 0.9 | 1.8 |
| | | A | 570 | 2.35 | 12.37 | | 93.5 [S:75.8,D:17.7] | 1.2 | 5.3 |

[ß] Estimated ploidy (EP). Refer to haplotig purging section in Methods.

[∂] Haplotigs were purged using the purge module within hifiasm (H. Cheng et al. 2020).

[Δ] gcpp-polished assembly was purged using purge_dups (Guan et al. 2020).

[ø] HiFi assembly was purged using purge_dups as hifiasm purging module didn't produce best results.

**Supplementary Table 6: BUSCO scores using the eudicotyledons_odb10 database with a total of 2121 conserved BUSCOs.**

| Species | Assembly type | Complete % | | Fragmented % | Missing % |
|---|---|---|---|---|---|
| | | Single % | Duplicate % | | |
| *Eucalyptus rudis* | HiFi | 96.6 | | 1 | 2.4 |
| | | 17.5 | 79.1 | | |
| | CLR | 97.1 | | 0.6 | 2.3 |
| | | 27.5 | 69.6 | | |
| *E. camaldulensis* | HiFi | 96.7 | | 0.8 | 2.5 |
| | | 13.9 | 82.8 | | |
| | CLR | 96.9 | | 0.7 | 2.4 |
| | | 13.7 | 83.2 | | |
