## Supplementary Figure 01 for "LeafGo: Leaf to Genome, a quick workflow to produce high-quality *De novo* genomes with Third Generation Sequencing technology"

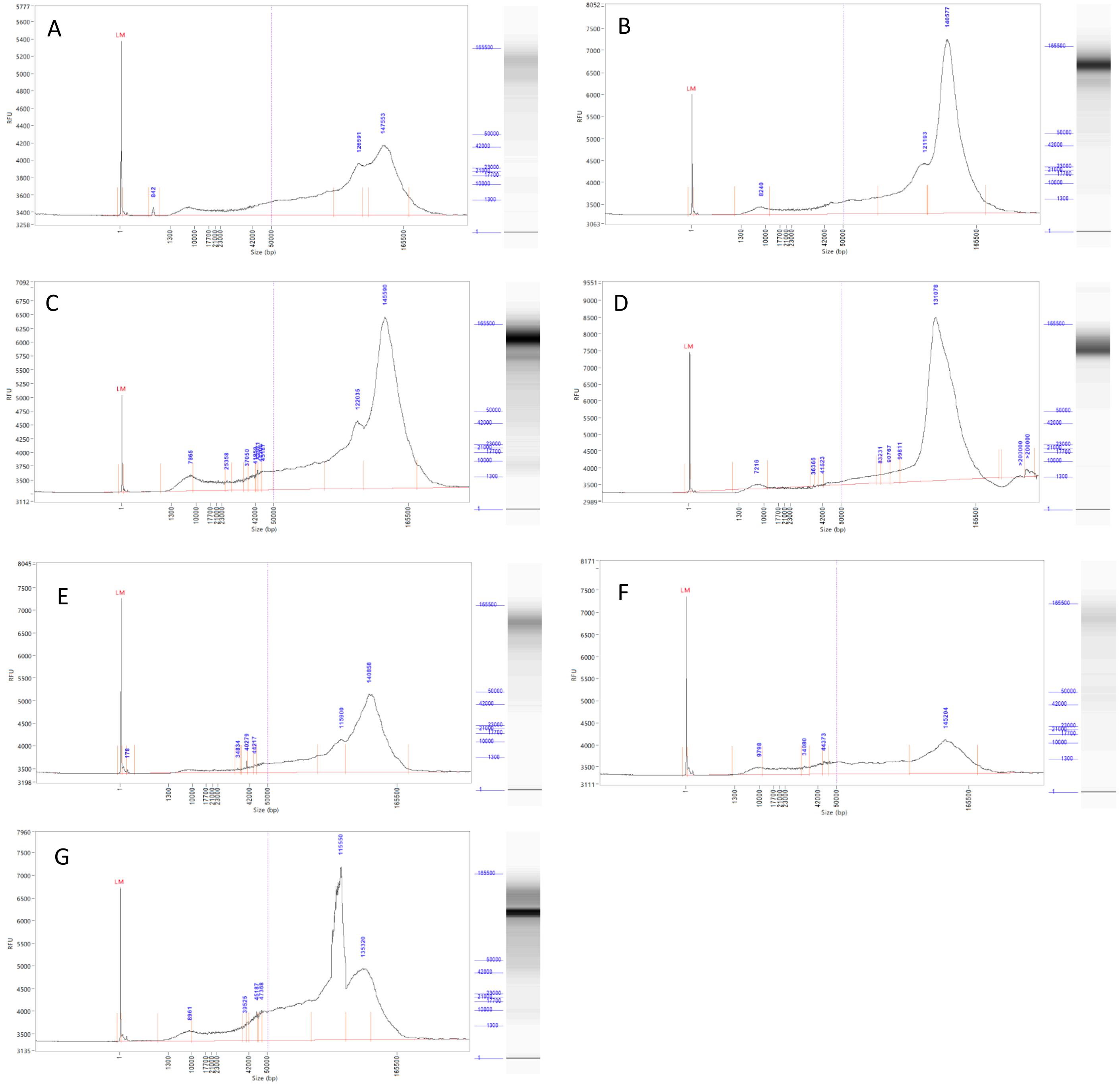
A
LM
842
126591
147553
RFU
Size (bp)
B
LM
8240
121193
140577
C
LM
7865
25358
37050
122035
145590
D
LM
7216
36366
41623
83231
90767
99811
131078
>200000
E
LM
178
34834
40279
44217
115900
140858
F
LM
9798
34080
44373
145204
G
LM
8961
39525
45187
47368
115550
135320

**Supplementary Figure 1: Capillary electrophoresis of the HMW DNA from seven plants.**

FEMTO 165 Kb ladder analysis (Method: FP-1002E22 – Extended gDNA 165 Kb) of extracted DNA from *Eucalyptus camaldulensis* subsp. *obtusa* (A); *E. rudis* (B); *Distichlis palmeri* (C); *Salvadora persica* (D); *Pennisetum glaucum* (E); *Zea mays* (F); and *Salicornia biglovii* (G).
