## Supplementary Figure 02 for "LeafGo: Leaf to Genome, a quick workflow to produce high-quality *De novo* genomes with Third Generation Sequencing technology"

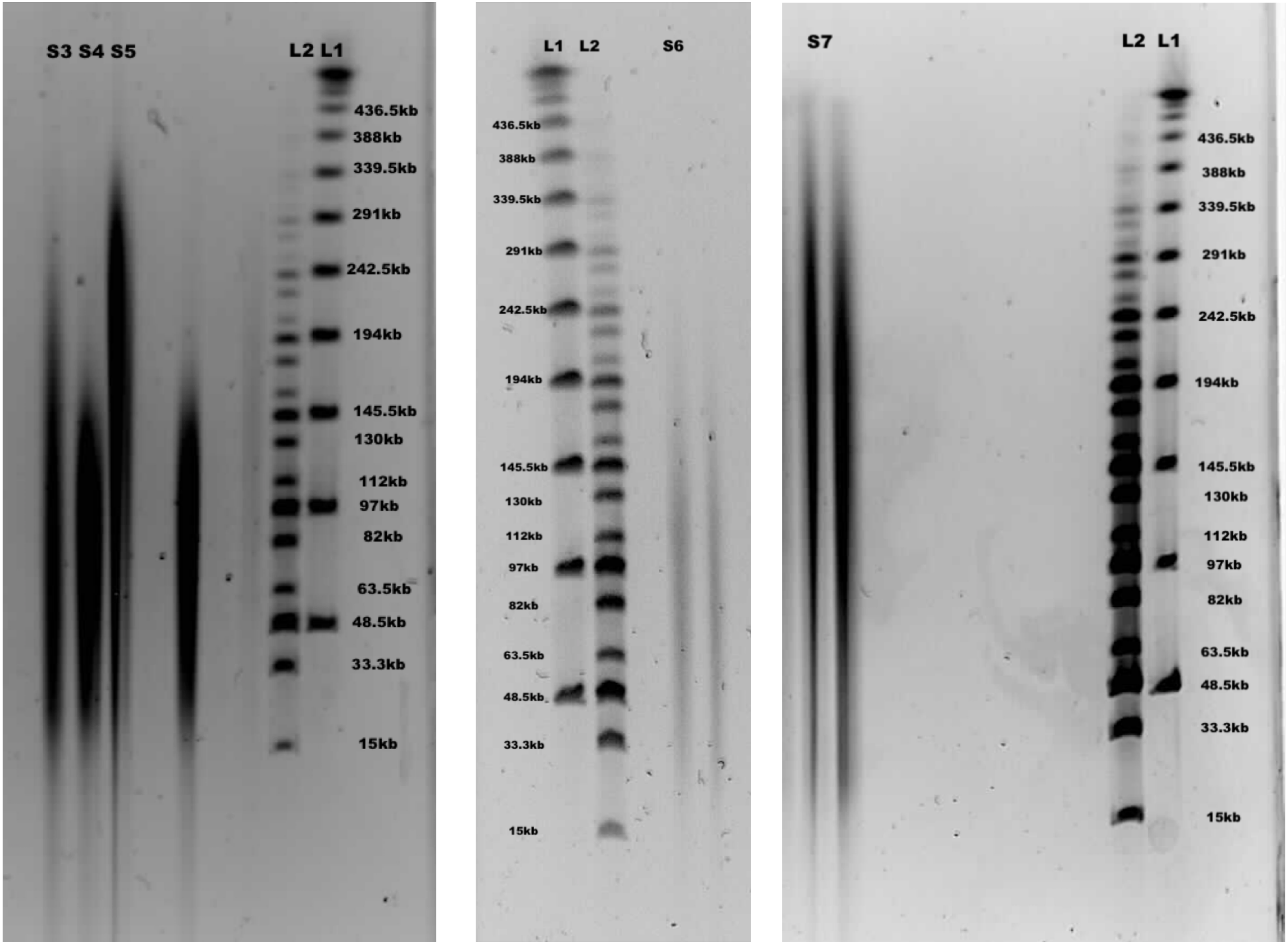
S3 S4 S5
L2 L1
436.5kb
388kb
339.5kb
291kb
242.5kb
194kb
145.5kb
130kb
112kb
97kb
82kb
63.5kb
48.5kb
33.3kb
15kb
L1 L2
S6
436.5kb
388kb
339.5kb
291kb
242.5kb
194kb
145.5kb
130kb
112kb
97kb
82kb
63.5kb
48.5kb
33.3kb
15kb
S7
L2 L1
436.5kb
388kb
339.5kb
291kb
242.5kb
194kb
145.5kb
130kb
112kb
97kb
82kb
63.5kb
48.5kb
33.3kb
15kb

**Supplementary Figure 2: Pulse field gel electrophoresis of extracted plant HMW DNA**

PFGE Image of HMW DNA, Lambda PFG Ladder (NEB N0341S) (L1), Midrange PFG Marker (NEB N0342S) (L2), Extracted DNA from *Eucalyptus rudis* (S1), *E. camaldulensis* (S2), *Zea Mays* (S3), *Distichlis palmeri* (S4), *Salvadora persica* (S5)*, Salicornia biglovii (S6),* and *Pennisetum glaucum* (S7).
