## Supplementary Figure 03 for "LeafGo: Leaf to Genome, a quick workflow to produce high-quality *De novo* genomes with Third Generation Sequencing technology"

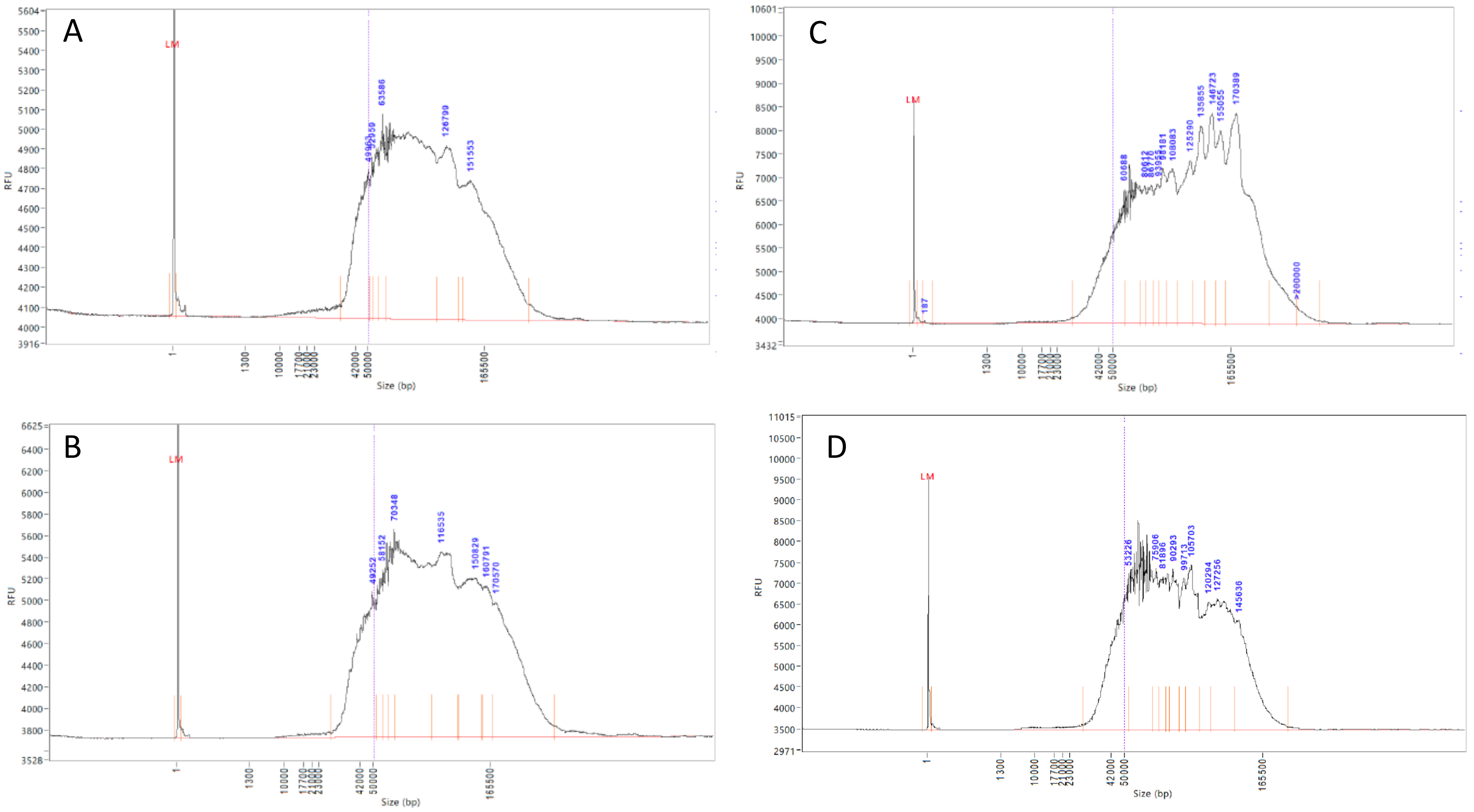
A
LM
63586
126790
151553
RFU
Size (bp)
1
1300
10000
17700
42000
50000
165500
B
LM
70348
116535
150829
160791
170570
49252
58152
C
LM
60688
108083
125290
135855
146723
155055
170389
187
>200000
D
LM
53226
75906
81896
90293
99713
105703
120294
127256
145636

**Supplementary Figure 3: Capillary electrophoresis of prepared CLR libraries**

FEMTO Pulse (Method: FP-1002E22 – Extended gDNA 165 Kb), results for PacBio CLR library of *Distichlis palmeri* (A), *Salvadora persica* (B), *Eucalyptus rudis* (C), *E. camaldulensis* (D).
