## Supplementary Figure 04 for "LeafGo: Leaf to Genome, a quick workflow to produce high-quality *De novo* genomes with Third Generation Sequencing technology"

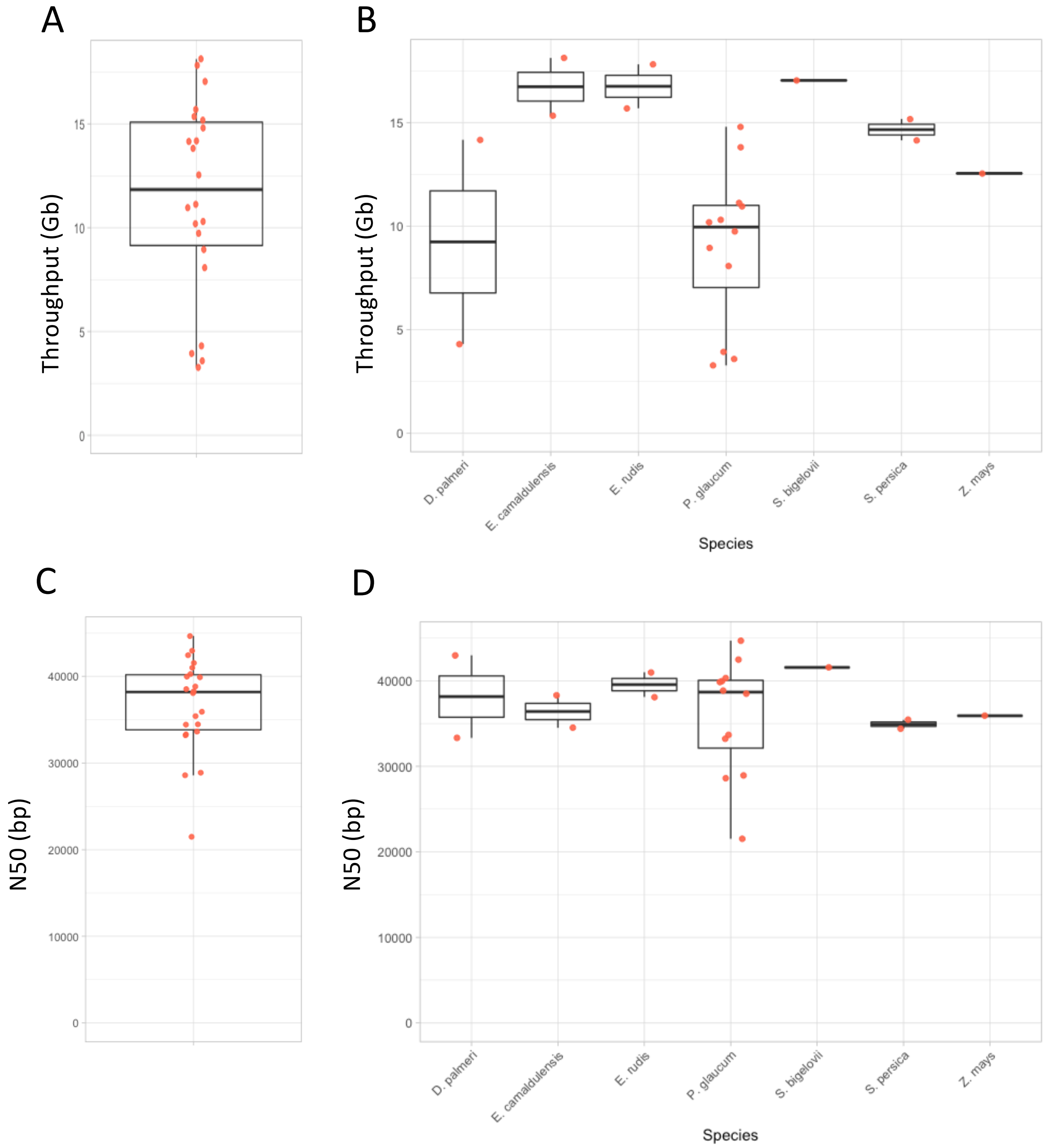

A
B
C
D
Throughput (Gb)
N50 (bp)
Species
D. palmeri
E. camaldulensis
E. rudis
P. glaucum
S. bigelovii
S. persica
Z. mays

**Supplementary Figure 4: Sequencing Throughput and N50 for PacBio CLR Libraries**

The throughput for all the sequenced libraries (A) and separated by species (B); the subread N50 for all the sequenced libraries (C) and separated by species (D). Sequencing platform: PacBio Sequel I.
