## Supplementary Figure 05 for "LeafGo: Leaf to Genome, a quick workflow to produce high-quality *De novo* genomes with Third Generation Sequencing technology"

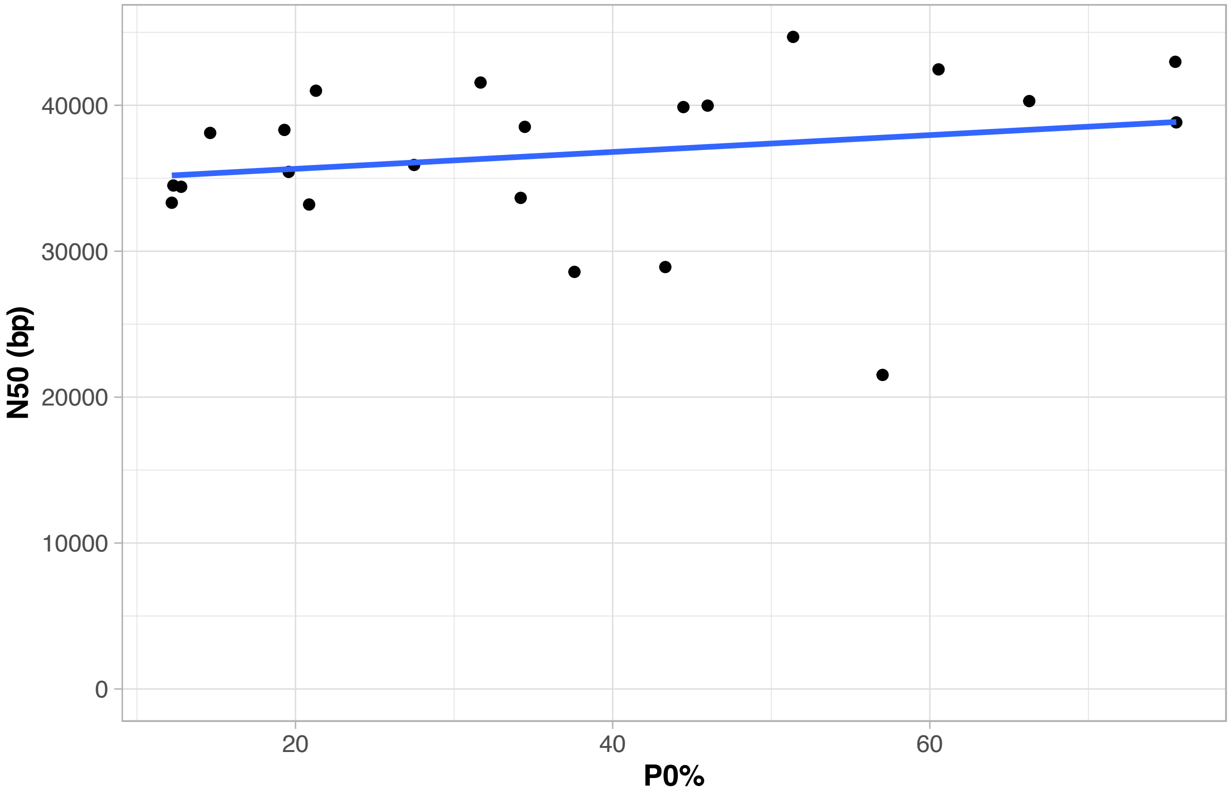

N50 (bp)
40000
30000
20000
10000
0
20
40
60
P0%

**Supplementary Figure 5: Correlation N50 vs P0%.**

Correlation between ZMW occupancy or library underloading (P0%) and subread N50 of CLR libraries sequenced with Sequel I. Spearman's rho = 0.42, p = 0.0499.
