## Supplementary Figure 06 for "LeafGo: Leaf to Genome, a quick workflow to produce high-quality *De novo* genomes with Third Generation Sequencing technology"

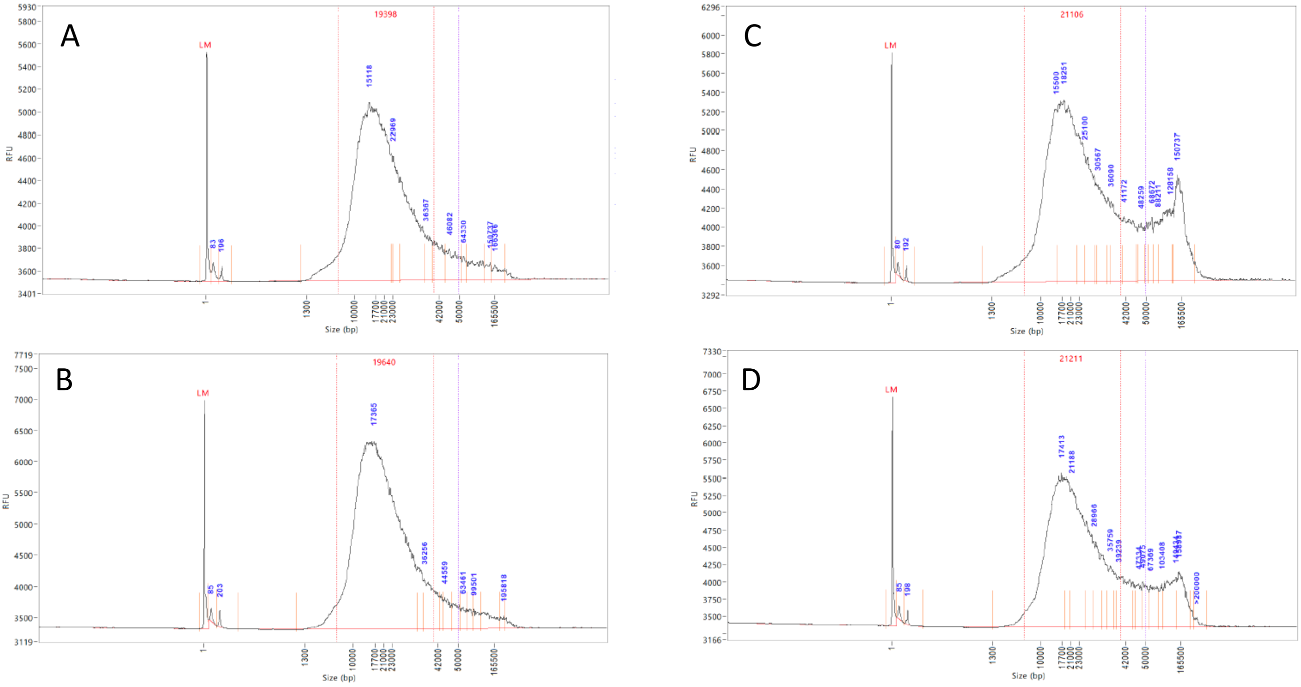
A
19398
LM
RFU
Size (bp)
15118
22969
36367
46082
64330
83
196
1
1300
10000
17700
21000
23000
42000
50000
165500
B
19640
LM
RFU
Size (bp)
17365
36256
44559
63461
99501
195818
85
203
C
21106
LM
RFU
Size (bp)
15500
18251
25100
30567
36090
41172
48259
68672
88211
128158
150737
80
192
D
21211
LM
RFU
Size (bp)
17413
21188
28966
35750
39239
103408
>200000
85
198

**Supplementary Figure 6: Capillary electrophoresis of prepared HiFi libraries**

FEMTO Pulse (Method: FP-1002-22 – gDNA 165 Kb), results for PacBio HiFi library of *Eucalyptus camaldulensis* before (A) and after size selection (B), and *E. rudis* before (C) and after size selection (D).
