## Supplementary Figure 07 for "LeafGo: Leaf to Genome, a quick workflow to produce high-quality *De novo* genomes with Third Generation Sequencing technology"

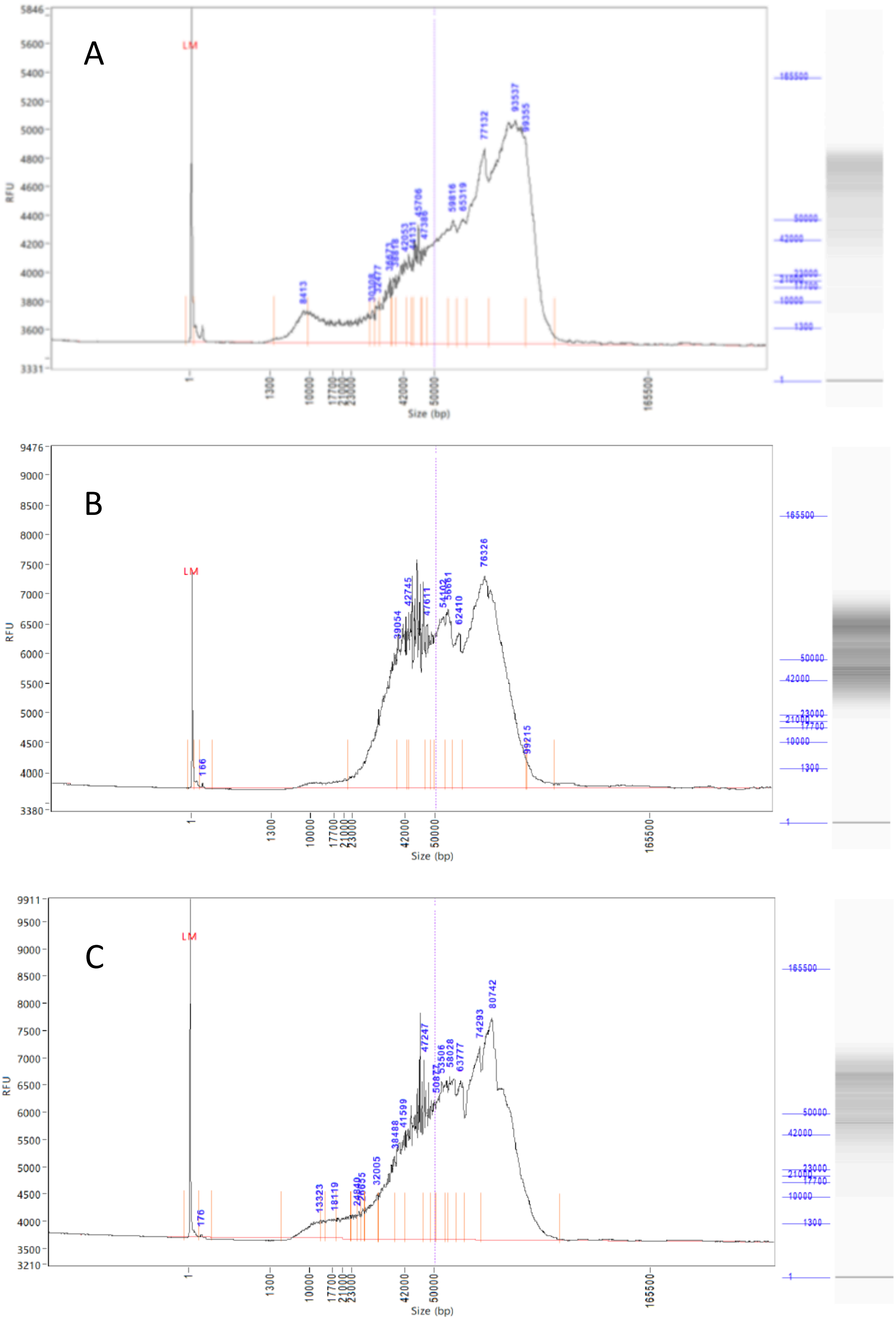
A
B
C
LM
RFU
Size (bp)
1
1300
10000
17700
21000
23000
42000
50000
165500
8413
77132
93537
76326
99215
166
176
80742
74293
47247
32005
13323
18119
41599
38488
62410
42745
39054

**Supplementary Figure 7: Size selection of ONT libraries**

Capillary electrophoresis (FEMTO Pulse, Method: FP-1002E22 – Extended gDNA 165kb) of gDNA (A), BluePippin 30 Kb selected DNA (B), and SRE-XL size selected DNA (C), from *Eucalyptus rudis* subsp. *rudis*.
