## Supplementary Figure 08 for "LeafGo: Leaf to Genome, a quick workflow to produce high-quality *De novo* genomes with Third Generation Sequencing technology"

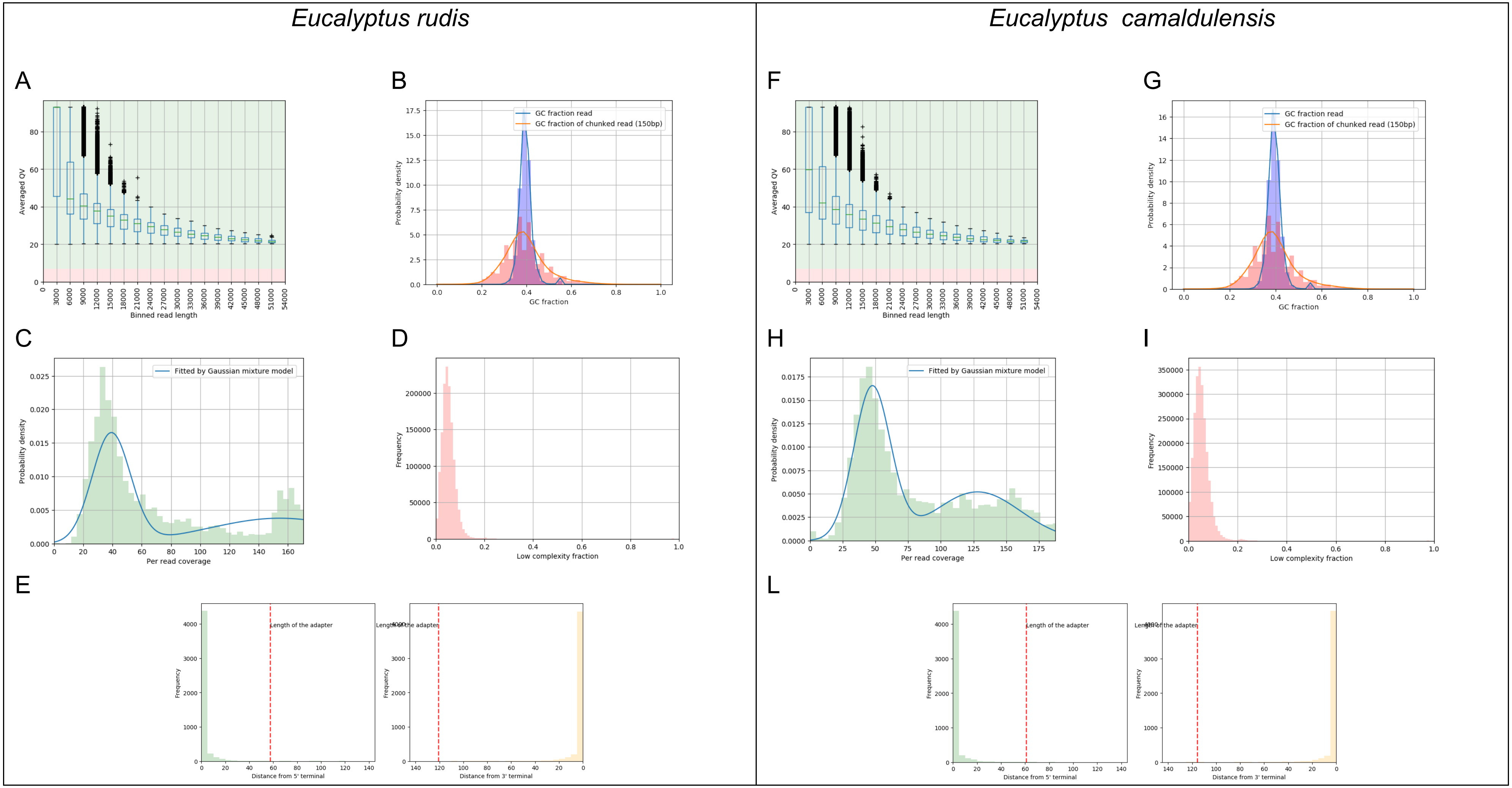

Eucalyptus rudis
Eucalyptus camaldulensis
A
B
C
D
E
F
G
H
I
L
Averaged QV
Binned read length
Probability density
GC fraction
GC fraction read
GC fraction of chunked read (150bp)
Fitted by Gaussian mixture model
Per read coverage
Frequency
Low complexity fraction
Length of the adapter
Distance from 5' terminal
Distance from 3' terminal

**Supplementary Figure 8: LongQC plots for HiFi data**

Sample data QC plots for PacBio HiFi dataset of *Eucalyptus rudis* and *E. camaldulensis* generated by LongQC (A, F) read-QV distribution alongside length (B, G) GC content, (C, H) estimated depth, (D, I) sequence complexity and (E, L) flanking region analysis.
