## Supplementary Figure 09 for "LeafGo: Leaf to Genome, a quick workflow to produce high-quality *De novo* genomes with Third Generation Sequencing technology"

### *Eucalyptus rudis*

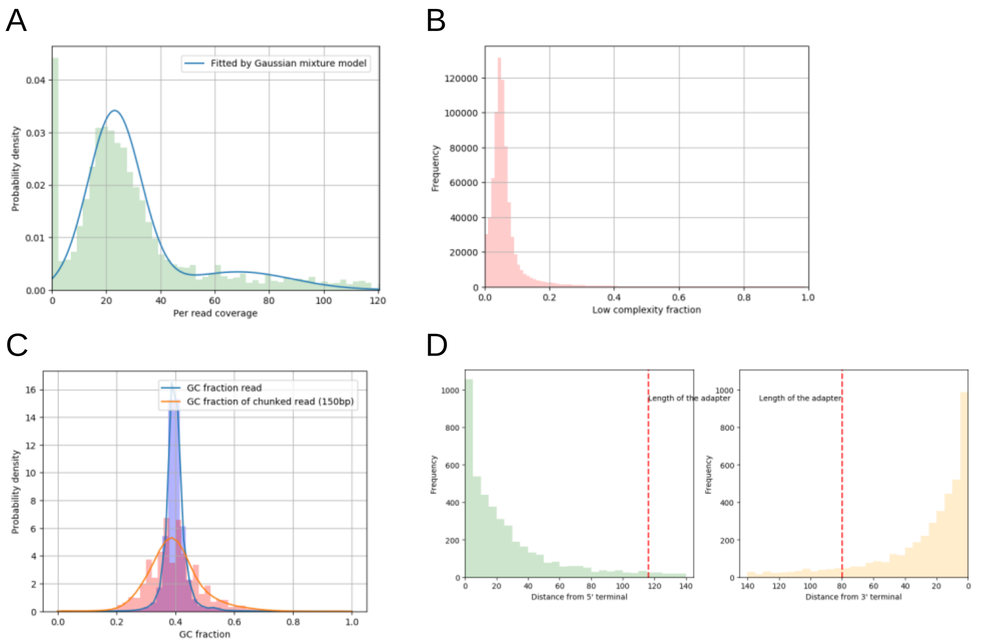


### *Eucalyptus camaldulensis*

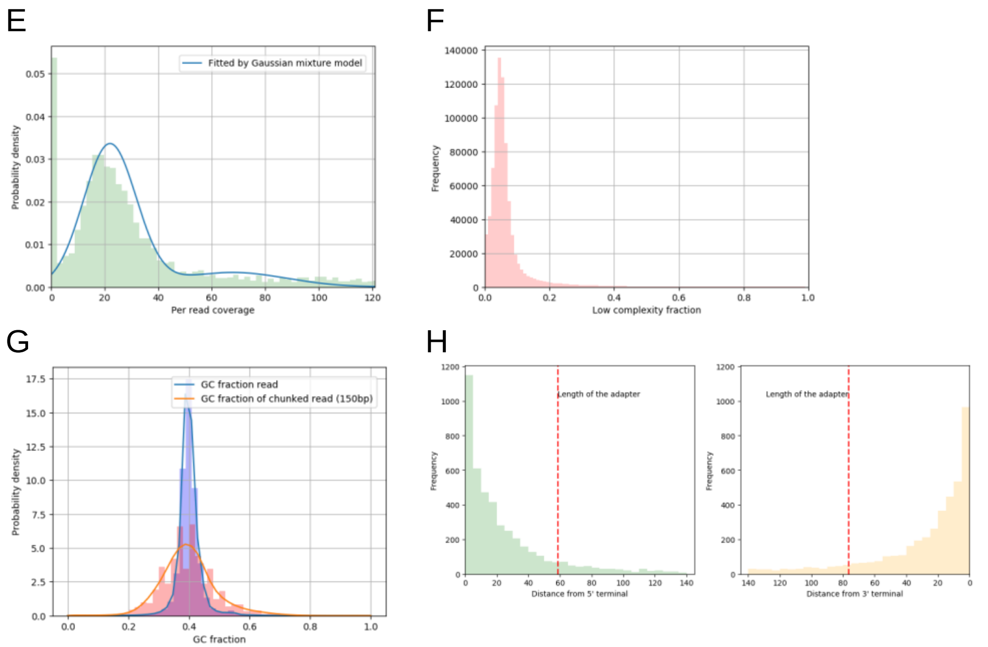

**Supplementary Figure 9: LongQC plots of CLR data for two *Eucalyptus* species**

Sample data QC plots for PacBio CLR dataset of *Eucalyptus rudis* and *E. camaldulensis* generated by LongQC (A,E) estimated depth (B,F) sequence complexity (C,G) GC content and (D,H) flanking region analysis.
