## Supplementary Figure 10 for "LeafGo: Leaf to Genome, a quick workflow to produce high-quality *De novo* genomes with Third Generation Sequencing technology"

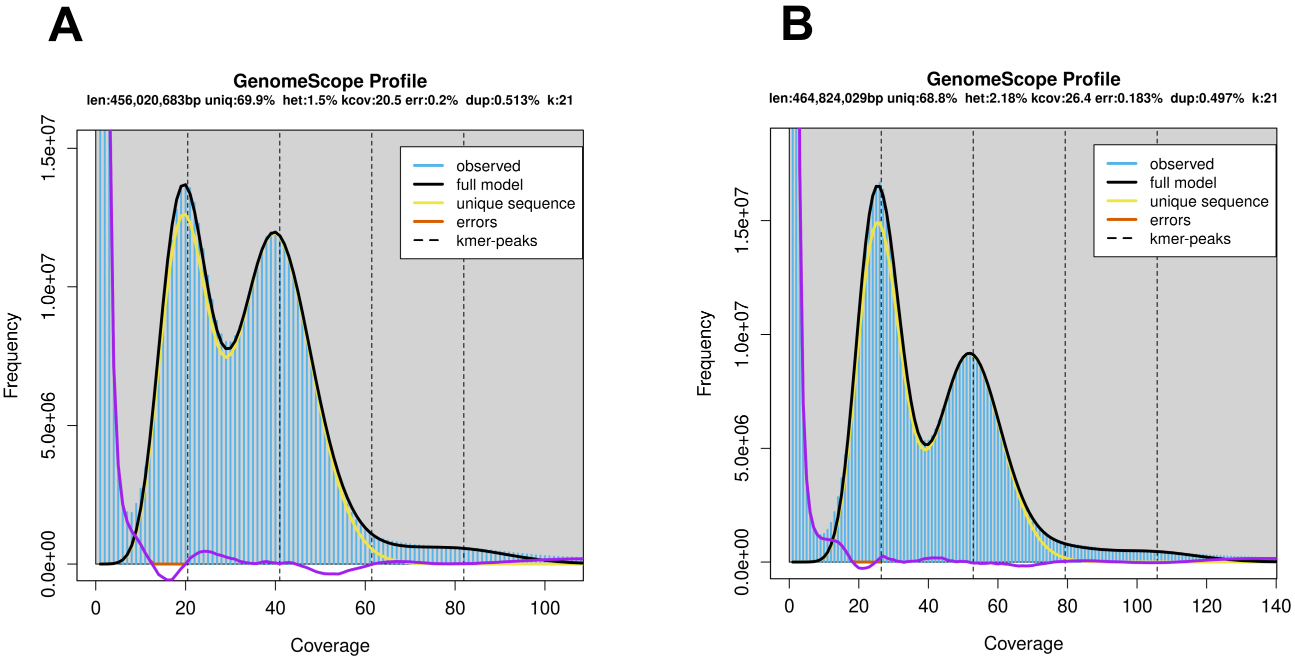
A
GenomeScope Profile
len:456,020,683bp uniq:69.9% het:1.5% kcov:20.5 err:0.2% dup:0.513% k:21
observed
full model
unique sequence
errors
kmer-peaks
Frequency
Coverage
0.0e+00
5.0e+06
1.0e+07
1.5e+07
0
20
40
60
80
100
B
GenomeScope Profile
len:464,824,029bp uniq:68.8% het:2.18% kcov:26.4 err:0.183% dup:0.497% k:21
observed
full model
unique sequence
errors
kmer-peaks
Frequency
Coverage
0.0e+00
5.0e+06
1.0e+07
1.5e+07
0
20
40
60
80
100
120
140

**Supplementary Figure 10: Genome profiling of the two *Eucalyptus* species based on HiFi data**

k-mer profiles, fitted models, and estimated parameters for the diploid genomes of *E. rudis* (A) and *E. camaldulensis* (B).
