## Supplementary Figure 11 for "LeafGo: Leaf to Genome, a quick workflow to produce high-quality *De novo* genomes with Third Generation Sequencing technology"

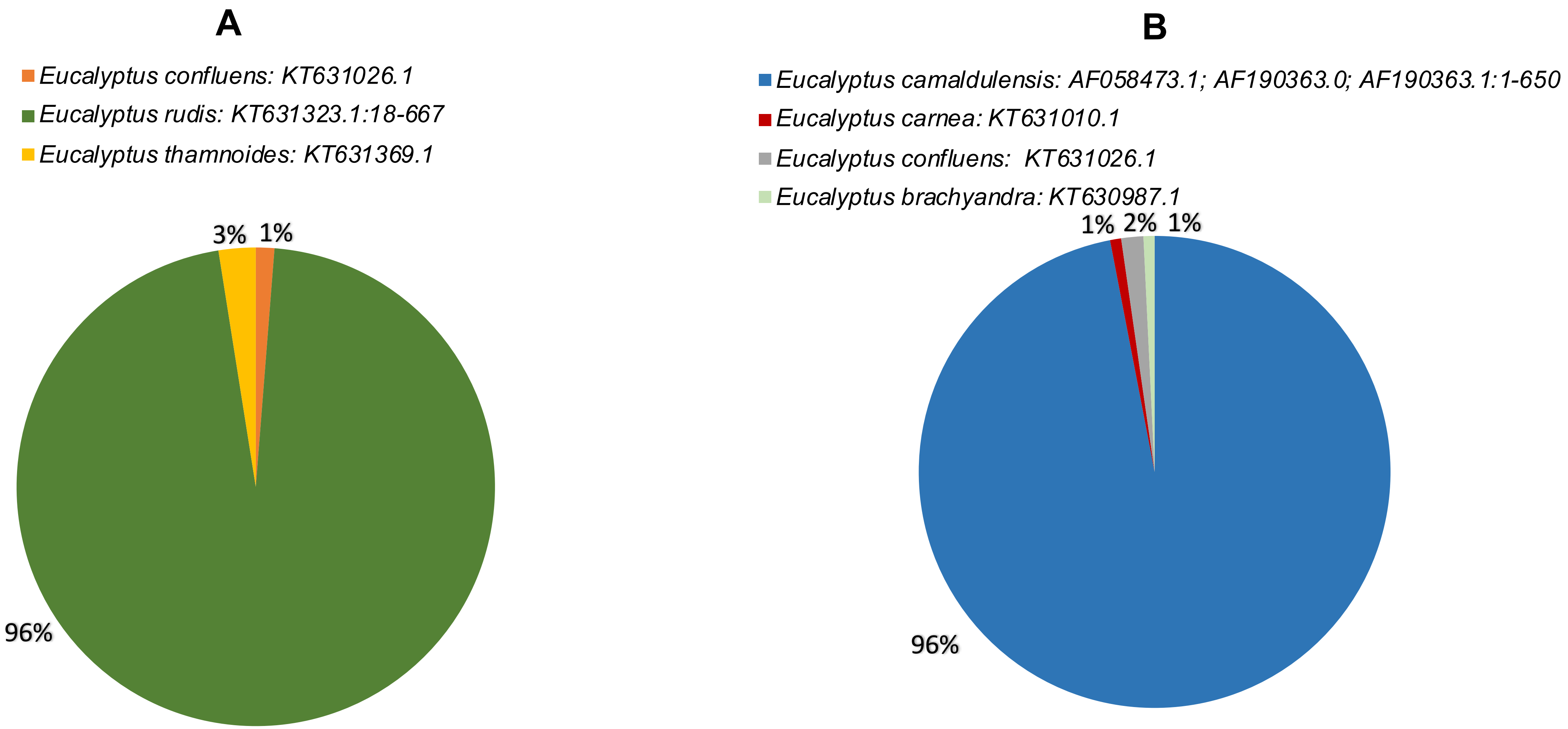
A
Eucalyptus confluens: KT631026.1
Eucalyptus rudis: KT631323.1:18-667
Eucalyptus thamnoides: KT631369.1
3%
1%
96%
B
Eucalyptus camaldulensis: AF058473.1; AF190363.0; AF190363.1:1-650
Eucalyptus carnea: KT631010.1
Eucalyptus confluens: KT631026.1
Eucalyptus brachyandra: KT630987.1
1%
2%
1%
96%

**Supplementary Figure 11: In silico Taxonomic classification of the two *Eucalyptus* species**

Identification of *Eucalyptus rudis* (A) and *E. camaldulensis* (B) by DNA metabarcoding analysis of both *Eucalyptus* genomes against the ITS dataset. Each pie chart shows the proportion of species identified by the top hit of each query.
